# Functional role of *pax6* in eye and central nervous system development in the annelid *Capitella teleta*

**DOI:** 10.1101/481135

**Authors:** Marleen Klann, Elaine C. Seaver

**Affiliations:** Whitney Laboratory for Marine Bioscience, University of Florida, 9505 Ocean Shore Blvd., St. Augustine, USA, Fl 32080; School of Life Sciences, University of Sussex, Brighton, United Kingdom, BN1 9RH

**Keywords:** annelid, *Capitella teleta*, pax6, eye, nervous system, morpholino, *pax6*, eye, nervous system, annelid, morpholino, *Capitella*

## Abstract

The transcription factor Pax6 is an important regulator of early animal development. Loss of function mutations of *pax6* in a range of animals results in a reduction or complete loss of the eye, a reduction of a subset of neurons, and defects in axon growth. There are no studies focusing on the role of *pax6* during development of any lophotrochozoan representative, however, expression of *pax6* in the developing eye and nervous system in a number of species suggest that *pax6* plays a highly conserved role in eye and nervous system formation. We investigated the functional role of *pax6* during development of the marine annelid *Capitella teleta*. Expression of *pax6* transcripts in *C. teleta* larvae is similar to patterns found in other animals, with distinct subdomains in the brain and ventral nerve cord as well as in the larval and adult eye. To perturb *pax6* function, two different splice-blocking morpholinos were used. Larvae resulting from injections with either morpholino show a reduction of the *pax6* transcript, and development of both the larval eyes and the central nervous system architecture are highly disrupted. Preliminary downstream target analysis confirms disruption in expression of some components of the retinal gene regulatory network, as well as disruption of genes involved in nervous system development. Results from this study, taken together with studies from other species, reveal an evolutionarily conserved role for *pax6* in eye development, and in neural specification and development.

## Introduction

*Pax* genes are a diverse family of transcription factors that function in a range of developmental processes (Mansouri, Hallonet, & Gruss, 1996; Noll, 1993), and are mainly characterized by the presence of two DNA binding domains, a paired domain and a paired-type homeodomain; each domain recognizes different downstream target genes (Chi & Epstein, 2002). The Pax6 transcription factor has two major roles during animal development. First, it functions early in an eye induction cascade to initiate transcription of numerous genes involved in eye development. In vertebrates, the paired-type homeodomain has been shown to be important for developmental processes of the eye such as lens formation and retinal specification (Ashery-Padan, Marquardt, Zhou, & Gruss, 2000; van Heyningen & Williamson, 2002). Second, Pax6 broadly regulates various other aspects of neurogenesis. For example, the balance between neural progenitor proliferation and differentiation during brain development is controlled by the paired domain (Haubst et al., 2004; Walcher et al., 2013). The importance of Pax6 during these two crucial developmental processes is also evident by identification of numerous target genes, which have been studied mainly in mouse (Holm et al., 2007; Purcell, Oliver, Mardon, Donner, & Maas, 2005; Sansom et al., 2009; Visel et al., 2007), zebrafish (Coutinho et al., 2011) and fruit fly (Halder et al., 1998; Ostrin et al., 2006). Direct and predicted targets such as *opsin* (Sheng, Thouvenot, Schmucker, Wilson, & Desplan, 1997), *eyes absent* (Coutinho et al., 2011; Halder et al., 1998; Purcell et al., 2005), *six1/2* (Halder et al., 1998; Ostrin et al., 2006), *six3/6* (Coutinho et al., 2011; Ostrin et al., 2006; Purcell et al., 2005), and *dachshund* (Coutinho et al., 2011; Purcell et al., 2005) are essential during eye formation. Likewise, target genes such as *neurogenin* (Bel-Vialar, Medevielle, & Pituello, 2007; Scardigli, Bäumer, Gruss, Guillemot, & Le Roux, 2003), *neuroD* (Coutinho et al., 2011; Holm et al., 2007; Sansom et al., 2009; Visel et al., 2007), *achaete-scute* (Holm et al., 2007; Sansom et al., 2009), *synaptotagmin IV* (Ostrin et al., 2006), and *pax2* (Coutinho et al., 2011; Matsunaga, Araki, & Nakamura, 2000) are indispensable for nervous system development. Moreover, in mouse and fly, homozygous *pax6* mutants are embryonic lethal, while heterozygous mutants have reduced or no eyes, and show underdeveloped brain structures and neural pathfinding defects (Halder et al., 1998; Hill et al., 1991; Huettl et al., 2016; Nomura, Haba, & Osumi, 2007).

There are numerous studies on the function of *pax6* in ecdysozoans and chordates (Aleen Remez et al., 2017; Ericson et al., 1997; Haubst et al., 2004; Huettl et al., 2016; Nakayama et al., 2015). However, although there is available expression data for a handful of species within lophotrochozoans, there is only one published article on the functional involvement of *pax6* in this superclade, and specifically demonstrates a role for *pax6* during eye regeneration in planarians (Pineda et al., 2002). Therefore, it is currently unknown if *pax6* has a central role in development of the eye and nervous system in this large and diverse animal clade. Although substantial progress has recently been made to develop techniques that alter gene expression, there are still many animal species where functional genomic studies are not yet established. The species-rich and diverse lophotrochozoan super phylum is one of the main three bilaterian clades, and is comprised of annelids, mollusks, platyhelminthes, and brachiopods among others. To date, only a few representatives within Lophotrochozoa have been successfully shown to be amenable for functional genomic studies, including the annelids *Platynereis dumerilii* (Bannister et al., 2014; Conzelmann et al., 2013) and *Helobdella robusta* (Song, Huang, Chang, & Weisblat, 2002), the mollusks, *Tritia obsoleta* (Rabinowitz, Chan, Kingsley, Duan, & Lambert, 2008) and *Crepidula fornicata* (Henry, Perry, & Martindale, 2010; Perry & Henry, 2015), and the planarian species *Dugesia japonica* and *Girardia tigrina* (Pineda et al., 2002). In recent years the marine annelid *Capitella teleta* has been successfully used for investigations related to evolutionary and developmental processes, and also for regeneration and environmental studies (Amiel, Henry, & Seaver, 2013; de Jong & Seaver, 2016; Meyer, Boyle, Martindale, & Seaver, 2010; Meyer, Carrillo-Baltodano, Moore, & Seaver, 2015; Pechenik, Berard, & Kerr, 2000; Pechenik et al., 2016; Seaver, 2016; Sur, Magie, Seaver, & Meyer, 2017). To further add to the importance of *C. teleta* as a representative annelid, functional genomic studies need to be established in this system. The ability to alter gene expression is of critical importance since this enables the establishment of a direct link between expression of particular genes and their specific functions, and gives insights into the consequences of loss or gain. In the present study, we investigated the role of *pax6* during the development of a lophotrochozoan representative, *Capitella teleta*, a marine annelid. Functional data was acquired via injections of splice-blocking antisense morpholinos (MO) into uncleaved embryos of *C. teleta*. Embryos were raised to larval stages and analyzed for phenotypic changes resulting from *pax6* transcript reduction.

## Materials and Methods

### Animal husbandry of *C. teleta* and *Ct-pax6* morpholino injections

A culture of *C. teleta* was kept in the laboratory following previously described methods (Seaver, Thamm, & Hill, 2005). Metamorphosis was induced by adding mud to competent larvae (Seaver et al., 2005). To obtain uncleaved eggs for morpholino injections, five females and four males were separated for at least two days, and then combined 11 hours before checking for presence of fertilized eggs. After dissecting the eggs from the brood tube, the egg membrane was softened by a 25 sec incubation with a freshly prepared 1:1 solution of 1M Sucrose:0.25M sodium citrate, with at least three subsequent washes in 0.2 μm filtered seawater (FSW). Quartz needles (QF 100-50-10, Sutter Instruments) were pulled on a micropipette puller (P-2000, Sutter Instruments) and used for pressure-injections. The needles were filled with a mixture comprised of the desired concentration of antisense morpholino oligos, nuclease-free water, and a 1:10 dilution of 20 mg/mL red dextran (Texas Red^®^, Molecular Probes™, dissolved in FSW). Injected and uninjected animals from the same brood were raised in FSW (with 60 μg/ml penicillin and 50 μg/ml streptomycin added) in separate dishes, and compared to determine the health of the brood. A brood was considered healthy if more than 90% of the uninjected animals developed normally. Two different splice-blocking morpholinos (designed by Gene Tools) directed against *C. teleta Ct-pax6* were tested: MO-1 (5’ AAGGGAAGAGGAGAGCCTACCTCTC 3’, Gene Tools) blocks splicing at the second exon-intron boundary and MO-2 (5’ ACTGACAGATTCATTAGTCTTACCT 3’, Gene Tools) blocks splicing at the fourth exonintron boundary. A generic standard control morpholino (5’ CCTCTTACCTCAGTTACAATTTATA 3’, Gene Tools) was employed to detect any potential nonspecific toxicity of the *Ct-pax6* morpholinos. The sequences of all morpholinos employed in this study were checked for potential off-target binding site in other genes by performing a Blast search against the publically available *Capitella* genome. All morpholinos were injected into uncleaved eggs, so that the entire embryo was affected by the reduction of *Ct-pax6* expression.

### Fixation, immunohistochemistry and whole mount *in situ* hybridization of *C. teleta* larvae and juveniles

Animals were raised to larval stages, and prior to fixation, were relaxed in 1:1 FSW:0.37M MgCl_2_ for 15 min (see (de Jong & Seaver, 2016) for recovery and pre-treatment of juveniles prior to fixation). *C. teleta* larvae and juveniles were fixed with 4% PFA in FSW for 30 min at room temperature (antibody staining) or overnight at 4 °C (whole mount *in situ* hybridization). Samples for antibody staining were subsequently washed with PBS and processed within 2-3 weeks. Samples for whole mount *in situ* hybridization were washed with PBS, then gradually transferred into 100% methanol and stored at −20 °C for extended periods.

DIG-labelled RNA probes were prepared for *Ct-pax6* (accession number: EY648731.1) (1255 bp) and *Ct-r-opsin1* (accession number: MG225382) (1057 bp) according to standard protocols using the T7 MEGAscript kit (Invitrogen). The colorimetric *in situ* protocol published previously (Seaver & Kaneshige, 2006; Seaver, Paulson, Irvine, & Martindale, 2001) was followed using a 1 ng/μL DIG-labeled RNA probe with a hybridization step for 72 h at 65 °C. Detection of the DIG-labeled RNA probe was either carried out using NBT/BCIP (Seaver et al., 2001) or fast red following the manufacturer protocol (SigmaFast™ Fast Red TR/Naphthol AS-MX Tablets). The color reaction was stopped after 2 h for both methods of detection.

Antibody labeling followed previously published protocols (Meyer et al., 2015). Primary antibodies and the concentrations used were: 1:10 mouse anti-22C10 (Developmental Studies Hybridoma Bank), 1:200 rabbit anti-5HT (Immunostar) and 1:400 mouse anti-acetylated α- tubulin (Sigma). Secondary antibodies and the concentrations used were: 1:500 donkey anti-mouse Alexa 546 (Invitrogen), 1:300 goat anti-rabbit Alexa 488 (Invitrogen) and 1:400 anti-mouse HRP (Jackson Immunoresearch). In *C. teleta*, anti-22C10 specifically labels sensory cells of the eyes (Yamaguchi & Seaver, 2013). Tyramide signal amplification of the 22C10 signal was used occasionally. After the immunohistochemistry protocol, samples were washed twice for 10 min in Tyramide buffer (2M NaCl, 0.1M Boric Acid, pH 8.5), and followed with a 10 min incubation in Development Solution (Tyramide buffer, 1:1000 IPBA 20mg/mL stock diluted in DMF, 1:10000 3% H_2_0_2_) with 1:1000 Rhodamine-conjugated tyramide (self-made, protocol adapted after (Hopman, Ramaekers, & Speel, 1998). *C. teleta* larvae possess segmental chaetae in the trunk that are highly auto-fluorescent and, based on their distinct morphology and position, are uniquely identifiable. All larvae were counterstained with 1:1000 Hoechst 33342 (Life Technologies) for 30 min in PBS, transferred into 70% glycerol/PBS and subsequently mounted onto microscope slides for imaging.

### RT-qPCR

Typically, 150-200 larvae (stage 6) were pooled for one biological replicate. The larvae were put in TRIzol (Invitrogen) and stored at −80 °C until further processing. RNA was extracted from the homogenised samples following a standard phenol/chloroform protocol. Prior to cDNA synthesis (SuperScript^®^ III First-Strand, Invitrogen), the RNA was treated for at least 1 h with DNAse (2 Units, TURBO DNA-free ™, Invitrogen). The RNA input for each cDNA synthesis reaction (total volume of 20 μL) was standardized to 400 ng. Control samples to check for the presence of residual genomic DNA contamination were made for each RNA sample by omitting the reverse transcriptase enzyme during the cDNA synthesis reaction (henceforth referred to as “-RT-control”). A Lightcycler 480 Real-time PCR instrument (Roche) was used for RT-qPCR reactions with a SYBR green kit (LightCycler^®^ 480 SYBR Green I Master, Roche). Each RT-qPCR reaction contained the following mix: 5 μL 2x SYBR green mix, 2.5 μL H_2_O, 1 μL 5 μM forward primer, 1μL 5μM reverse primer, 0.5 μL cDNA. Technical triplicates were run for each individual biological replicate. Each experimental sample is represented by at least 3 biological replicates (injection into a set of embryos from different broods). PCR cycling conditions followed standard programs with 40x [10 sec 95 °C, 20 sec 62 °C, 20 sec 72 °C], except for the exon-exon primer pairs (see below) with 40x [20 sec 95 °C, 30 sec 62 °C, 30 sec 72 °C] due to an increased length of the expected fragment. To select a suitable reference gene for calculating relative expression levels, four house-keeping genes were tested: *Ct-CDC5* (*C. teleta cell division cycle 5*), *Ct-RPS9* (*C. teleta ribosomal protein S9*), *Ct-HPRT* (*C. teleta hypoxanthine phosphoribosyl transferase*), and *Ct-EF1* (*C. teleta elongation factor 1*) (Suppl. Fig. 1). *Ct- HPRT* was chosen as an appropriate reference gene for two main reasons. First, since this is a developmental study, it was important that the reference gene shows a rather stable expression across different developmental stages. Five larval stages, stage 4, stage 5, stage 6, stage 7, and stage 7/8 were examined (Suppl. Fig. 1). Since *Ct-CDC5* was found to have higher expression during larval stage 7/8, it was excluded. Second, the expression levels of *Ct-HPRT* (green curve in Suppl. Fig. 1) are most similar to the expression level of the gene of interest, *Ct-pax6* (blue curve in Suppl. Fig. 1), while *Ct-RPS9* and *Ct-EF1* expression levels are much lower (red and purple curves in Suppl. Fig. 1). Two sets of primers to validate the efficiency of the *Ct-pax6* morpholinos were designed for each of the two MOs. The sequences of all primers used can be found in Suppl. Table 1. The *Ct-pax6* primer sets whose targets are located on exons, F1&R1 and F3&R3, will detect a reduction of the spliced *Ct-pax6* transcript in larvae injected with MO, since the inclusion of the intron will make the qPCR amplicon too long to be synthesized. The *Ct-pax6* primer sets that are targeted to introns, F2&R2 and F4&R4, will detect unspliced *Ct- pax6* transcript only, meaning they will only amplify a product either in cDNA samples which have been modified due to MO activity, or in genomic DNA. cDNA samples used for analysis of potential downstream targets were the same as used for determination of the MO efficiency.

**Fig. 1.**
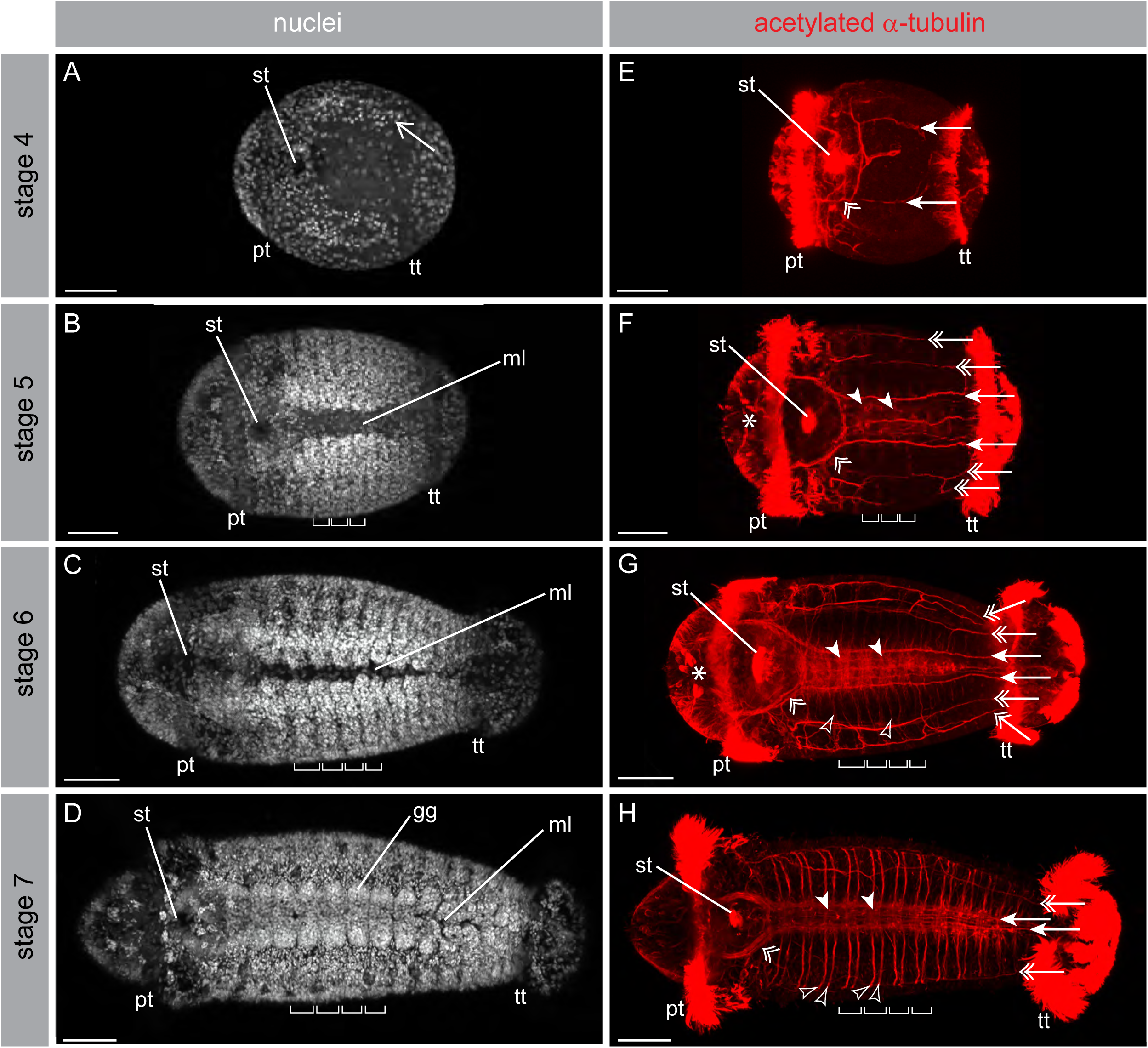
Characteristics of *C. teleta* larvae. Hoechst staining to label nuclei in white (nuclei; A-D) and anti-acetylated α-tubulin staining to label axon projections and cilia in red (a-tub^+^ NS; E- H). The images in each row are from the same individual, and all images are ventral views with anterior to the left. Four different larval stages are shown: stage 4 (A, E), stage 5 (B, F), stage 6 (C, G), and stage 7 (D, H). All images are z-stack projections, with anterior to the left. The open arrow in A shows the location of the segment anlagen. The brackets in B, C, D, F, G and H represent the width of individual segments. The double open arrowheads in E–H point towards the circumoral nerves. The main connectives of the ventral nerve cord are indicated by the arrows in E-H, while double open arrows point towards the lateral longitudinal nerves. In F-H, filled arrowheads indicate two sets of commissures in the ventral nerve cord. Unfilled arrowheads highlight selected individual segmental nerves in F-H. The asterisks in F and G mark the position of the anti-acetylated α-tubulin positive neurons in the head. Abbreviations: gg: ganglia, ml: ventral midline, pt: prototroch, st: stomodeum, tt: telotroch. Scale bars: 50 µm.

### Statistical analyses

Relative expression ratios were calculated using the method reported in Pfaffl (2001). This method takes into consideration that amplification efficiency between the reference gene and the experimental genes often is not exactly the same, and therefore primer efficiencies for individual genes are incorporated into the formula. Calculation of primer efficiency (E) and other statistical analyses were performed in Microsoft Excel. To calculate the primer efficiency, a standard curve was generated for each primer set for every individual qPCR reaction performed using the following dilutions of cDNA: undiluted, 1:2, 1:4, 1:8, 1:16 and 1:32. The average Ct values of technical triplicates were plotted, and a line of best fit was added to determine the slope. The primer efficiency is = 10^-^1^/*slope*^. This primer efficiency was taken into account when calculating the relative fold change values. Fold change was calculated as

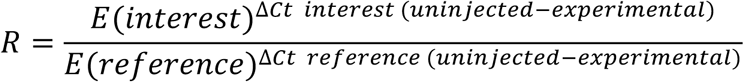

Student’s *t*-tests were used to determine if different treatment groups showed statistically significant relative fold changes (statistical significance if *p* ≤ 0.05). A two-tailed Student’s *t*-test was used to determine if there were statistically significant differences in the *Ct-pax6* transcripts of larvae resulting from control MO injections compared to larvae resulting from uninjected embryos. A one-tailed Student’s *t*-test was used to determine if there were statistically significant differences in the relative *Ct-pax6* transcript levels from larvae resulting from *Ct-pax6* MO injections compared to larvae resulting from control MO injections. For analysis of downstream target genes, a one-tailed Student’s t-test was used to calculate *p* values because the morphological phenotypic changes negatively affected the formation of the nervous system, eye and segment boundary. RT-qPCR results for one of the four biological replicates for the downstream target analysis varied quite a lot from the remaining three biological replicates. However, since multiple technical replicates of this particular biological replicate were consistent, this appears to represent biological variation. Therefore, all four biological replicates were pooled in order to calculate the final fold change values.

### Documentation and Analysis

Colorimetric *in situ* hybridization signals were documented with a digital Spot FLEX camera (Diagnostic Instruments) connected to an Axioskop2 mot plus compound microscope (Zeiss). Fluorescent labeling, including fast red *in situ* hybridization signal, antibody and nuclear staining, was documented with a Zeiss confocal laser scanning microscope 710 (Zeiss) using the ZEN 2011 software. In order to combine/render image stacks of individual larvae or juveniles taken with the Axioskop2 compound microscope, the software Helicon Focus (d-Studio Ltd.) was used (method B, radium 50 and smoothing 1-7). Confocal image stacks were analyzed and imaged with the 3D-reconstruction software IMARIS (Bitplane AG). Images were further processed in Adobe Photoshop CS3. Figure plates and schematic representations were created with Adobe Illustrator CS3.

## Results

### Overview of developmental characteristics in *C. teleta*

A detailed staging system of *C. teleta* has been published previously (Seaver et al., 2005) and will only be briefly summarized here. The early spiral cleavages represent stage 1, followed by late cleavage and gastrulation that are designated stage 2 and 3, respectively. At stage 4, about 2 days after fertilization, larvae start to show the first morphologically distinct features (Fig. 1A, E). The following developmental stages last about 1 day each: 3 days of development is stage 5 (Fig. 1B, F), 4 days of development is stage 6 (Fig. 1C, G), and 5 days of development is stage 7 (Fig. 1D, H). At stage 9, about 7-8 days after fertilization, larval development is complete and metamorphosis can be induced. In larvae with labeled nuclei (Fig. 1A-D), the prototroch (pt), telotroch (tt) and stomodeum (st) appear as regions of lower nuclear density. The prototroch and telotroch are ciliary bands that run around the circumference of the larvae just posterior of the head and anterior of the pygidium, respectively, and these prominent features are useful landmarks. Another landmark is the stomodeum, which also possesses cilia. The cilia are heavily labeled with anti-acetylated α-tubulin (Fig. 1E-H) and therefore all landmarks are clearly visible. Stage 4 larvae show lateral segment anlagen (Fig. 1A, open arrow) that will form the trunk segments and become visible in stage 5 larvae (Fig. 1B-D, brackets). Segmentation starts in the ventral lateral region (Seaver et al., 2005), leaving an unsegmented ventral midline (ml). Over time the segments expand dorsally and ventrally, and the unsegmented ventral midline successively gets more restricted (compare Fig. 1B, C and D). After complete closure of the ventral midline in the anterior trunk at stage 7, ganglia become visible (Fig. 1D, gg). A detailed description of the development and architecture of the nervous system in *C. teleta* has been published (Meyer et al., 2015). The first distinct anlagen of the central nervous system, the circumoral nerves (Fig. 1E-H, double open arrowheads) and main longitudinal connectives (Fig. 1E-H, arrows) appear at stage 4. Further maturation of the ventral nerve cord includes the addition of commissures at stage 5 (Fig. 1F-H, filled arrowheads) and segmental nerves at stage 6 (Fig. 1G, H, unfilled arrowheads). The latter are characteristic components of the peripheral nervous system, as are the lateral longitudinal nerves that run along the trunk, two on each side (Fig. 1F-H, double open arrows). Anterior to the brain is a bilateral cluster of anti-acetylated α- tubulin positive neurons, comprising three neurons on each side (Fig. 1F, G, asterisks).

The eyes in *C. teleta* are rather simple, and are comprised of only three different cell types: a pigment, sensory, and supporting cell (Rhode, 1993). The antibody 22C10 specifically labels sensory cells associated with the eyes in *C. teleta* (magenta in Fig. 2A’, A’’, B’, B’’, C’, C’’). Previous studies have demonstrated that the sensory neuron of the eye is distinctively labeled by 22C10 in annelids, such as *C. teleta* (Yamaguchi & Seaver, 2013) and *Platynereis massiliensis* (Helm, Adamo, Hourdez, & Bleidorn, 2014). The head region of older *Capitella* larvae (starting at late stage 5) has two bilateral 22C10 positive domains on each side of the head, an anterior sensory cell (magenta in Fig. 2A’, A’’, B’, B’’, C’, C’’, unfilled arrowheads) and a posterior sensory cell (magenta in Fig. 2A’, A’’, B’, B’’, C’, C’’, filled arrowheads). The anterior sensory cell (Fig. 2A, B, C, unfilled arrowheads) is situated medially within the brain (Fig. 2C, yellow outline), while the posterior sensory cell (Fig. 2A, B, C, filled arrowheads) is located towards the lateral-posterior edge of the brain. Double labeling experiments with *Ct-r-opsin1* (yellow in Fig. 2A’, A’’’, B’, B’’’) and the 22C10 antibody (magenta in Fig. 2A’, A’’, B’, B’’), clearly show colocalization (Fig. 2A’, B’), indicating the specificity of anti-22C10 to label photo-sensory cells in *C. teleta*. Fine neural processes from both the anterior and posterior sensory cell extend medially and appear to meet each other to form a single projection, which ends in an agglomeration straddling the midline of the brain (Fig. 2C-C”’; Fig. 9E, arrows). Comparison with antiserotonin labeling (green in Fig. 2C’, C’’’) reveals that the 22C10-positive projections are within the main cerebral commissure of the brain. This is in contrast to the situation found in the annelid *Platynereis dumerilii*, where there is a direct axonal contact between the eyespots and the prototroch (Jékely et al., 2008). The posterior sensory cell is in contact with the pigment cell of the larval eye (Yamaguchi & Seaver, 2013) (Fig. 9A, double open arrowheads), and the pigment cell is situated close to the prototroch at the lateral-posterior portion of the brain. The anterior 22C10-labeled cell is thought to be the progenitor of the juvenile eye as the posterior cell is lost at metamorphosis (Yamaguchi & Seaver, (2013).

**Fig. 2.**
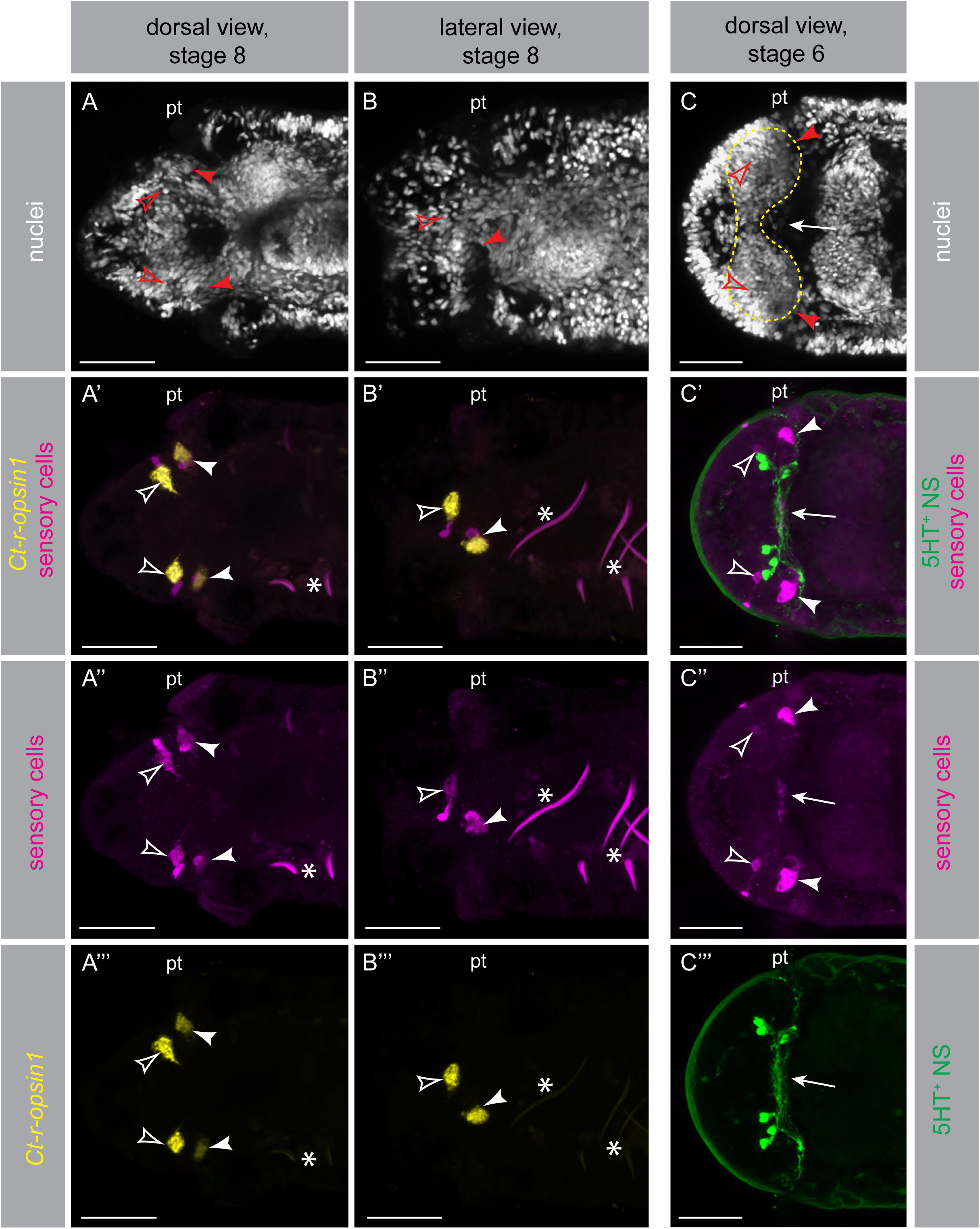
Characteristics of the eyes in *C. teleta* larvae. Hoechst staining to label nuclei in white (nuclei; A, B, C), *Ct-r-opsin1* whole mount *in situ* expression in yellow (*Ct-r-opsin1*; A’, A’’’, B’, B’’’), 22C10 antibody labeling of the sensory cell of the eye in magenta (sensory cells; A’, A’’, B’, B’’, C’, C’’), and serotonin antibody labeling of neural projections in green (5HT^+^ NS; C’, C’’’). A’, B’ and C’ are merged channels of A’’ plus A’’’, B’’ plus B’’’, and C’’ plus C’’’, respectively. The images shown are z-stack projections of the head-region, with anterior to the left. Images from each column are from the same individual. There are two photo-sensory cells, an anterior sensory cell (unfilled arrowheads) and a posterior sensory cell (filled arrowheads). The arrow marks the region where the neural projections of the sensory cells meet (C-C’’’). Segmentally iterated chaetae of the trunk are autofluorescent and therefore visible in some of the 22C10 images (asterisks). Abbreviation: pt: prototroch. Scale bars: 30 µm.

### *Ct-pax6* is expressed in distinct subdomains of the nervous system and the eye

A previous paper identified one *pax6* orthologue in the *C. teleta* genome (Seaver, Yamaguchi, Richards, & Meyer, 2012), which encodes two highly conserved DNA-binding domains, a bipartite paired domain and a paired-type homeodomain, and a C-terminal proline-serine-threonine (PST)-rich transactivation domain. A recent study of genome wide searches for homeodomain-containing genes in annelids reported a second *pax6* gene in the genome of *C. teleta* (Zwarycz, Nossa, Putnam, & Ryan, 2015), which is in line with other recent findings in several non-mammalian vertebrate species (Ravi et al., 2013). However, this putative second *pax6* gene, *pax6.2*, encodes only a Pax6-like homeodomain, but not a paired domain characteristic for genes belonging to the *pax* family. Multiple attempts to amplify this second *pax6* gene from various cDNA templates (different stages of development) were unsuccessful, and therefore this putative second *pax6* gene does not appear to be transcribed. From this observation, together with the fact that the predicted protein lacks a paired domain, we deduce that *C. teleta* has a single functional *pax6* ortholog, which we denote as *Ct-pax6*.

The wildtype expression pattern of *Ct-pax6* in *C. teleta* was investigated using *in situ* hybridization (Fig. 3, 4). Early cleavage and gastrulation stages did not show any detectable *Ct- pax6* signal (data not shown). The earliest detectable *Ct-pax6* expression is at larval stage 3, where the transcript is found in two small bilateral domains in the anterior ectoderm (data not shown). In early larval stage 4, the main expression domain of *Ct-pax6* is in the developing brain lobes (Fig. 3A, asterisk). In addition, there are a few scattered *Ct-pax6*-positive cells in the anlagen of the ventral nerve cord (Fig. 3A, filled arrowheads). This onset of *Ct-pax6* expression correlates with the appearance of differentiated neurons in the brain and the main nerves of the central nervous system, including a nerve connecting the two brain lobes, the circumoral nerves and the main longitudinal connectives (see Fig. 1E and (Meyer et al., 2015). By larval stage 5, *Ct-pax6* expression in the brain lobes has broadened medially and posteriorly (Fig. 3B, asterisk). The ventral nerve cord now exhibits segmentally iterated stripes of expression (Fig. 3B, black arrows). Brain expression in stage 6 larvae starts to segregate into a lateral (Fig. 3C, asterisk) and a medial (Fig. 3C, double open arrowhead) subdomain. Even more cells of the ventral nerve cord express *Ct-pax6* and expression spans the length of the trunk. Nevertheless, segmental stripes are clearly visible in the anterior four to five segments (Fig. 3C’, black arrows), while the expression in the posterior is denser. This pattern likely reflects the anterior to posterior temporal progression of ganglia formation in the ventral nerve cord. There is a clear distinction of the *Ct- pax6* brain expression into subdomains at larval stage 7, a lateral (Fig. 3D, asterisk), a medial (Fig. 3D, double open arrowhead) and a ventral expression domain (Fig. 3D, double open arrow). Expression in discrete subsets of cells in the ventral nerve cord has become more pronounced, displaying a segmentally iterated pattern of short lateral, diagonal stripes (Fig. 3D’, black arrows) and medial spots (Fig. 3D’, white arrows) in the anterior segments. Apart from the predominate expression of *Ct-pax6* in elements of the central nervous system, few isolated cells in the posterior 2/3 of the trunk ectoderm (Fig. 3D’, unfilled arrowheads) also express the transcript. Although elements of the peripheral nervous system have not been mapped carefully in *C. teleta*, the position of these *pax6*-positive cells might indicate that they are a part of the peripheral nervous system. A number of neurogenic genes, such as *prospero*, *soxB1*, and *soxB* have been shown to have similar expression patterns (Sur et al., 2017), supporting the idea that the peripheral nervous system develops in this trunk region. After the prominent expression of *Ct-pax6* at stage 6 and 7, transcription is down regulated at larval stage 8 (Fig. 3E, E’). This holds true for the bipartite brain expression (Fig. 3E, asterisk and double open arrowhead) as well as the ventral nerve cord where expression is most pronounced in the lateral portion of the ganglia (Fig. 3E’, filled arrowheads), with a few scattered cells in more medial positions. At larval stage 9, *Ct-pax6* expression is reduced further, with very small lateral and medial subdomains in the brain, and almost no expression in the ventral nerve cord (data not shown). Juvenile worms show a similar pattern of *Ct-pax6* expression in the brain as the larvae, a lateral (Fig. 3F, asterisk) and a medial (Fig. 3F, double open arrowhead) subdomain. Isolated, segmentally repeated cells within the ganglia of the ventral nerve cord are *Ct-pax6*-positive in juveniles (Fig. 3F, filled arrowheads).

**Fig. 3.**
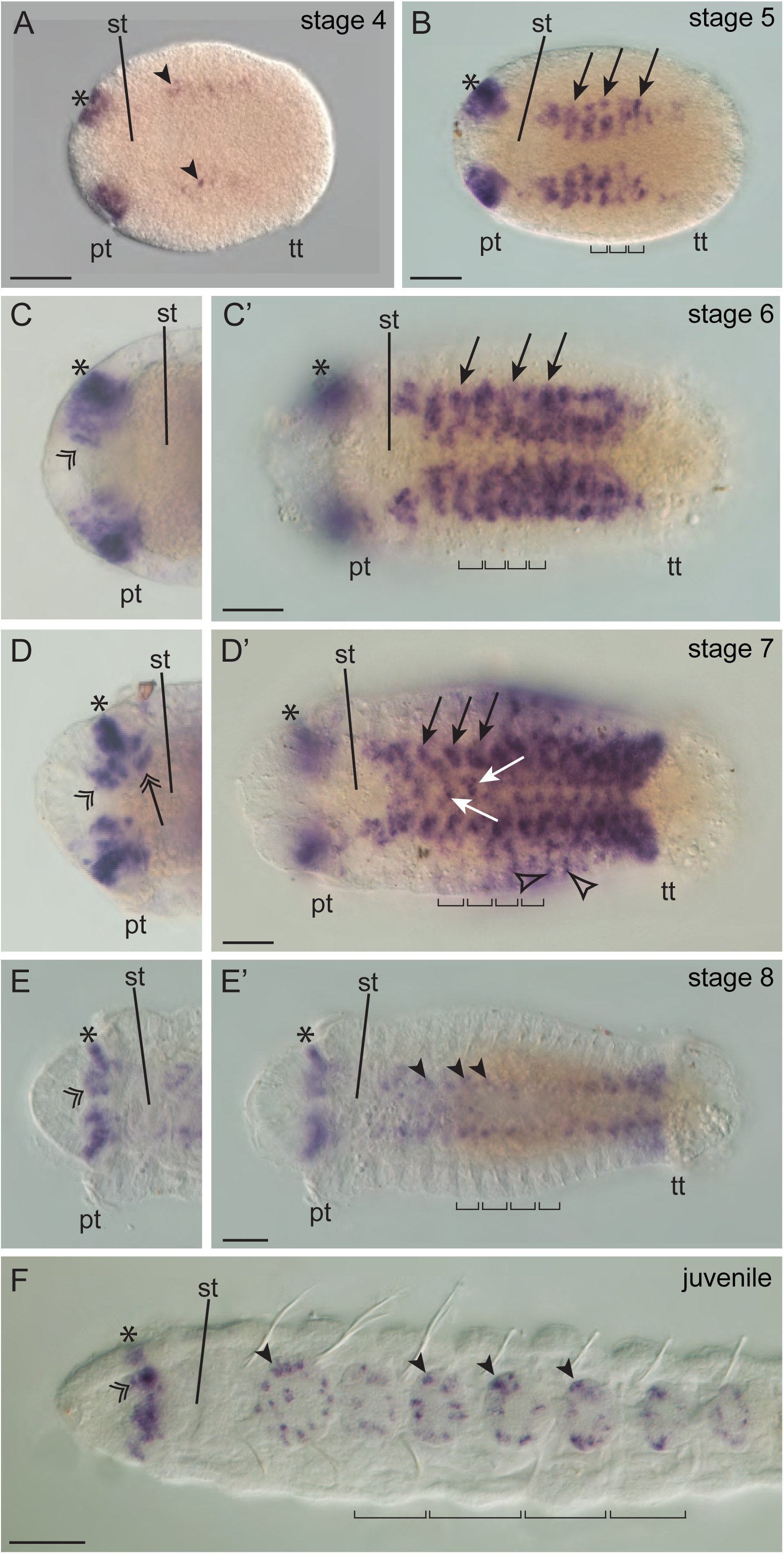
Expression of *Ct-pax6* in the nervous system. Colorimetric whole mount *in situ* for *Ct- pax6* (dark purple). Six different stages are shown: larval stage 4 (A), larval stage 5 (B), larval stage 6 (C, C’), larval stage 7 (D, D’), larval stage 8 (E, E’), and 12 days juvenile (F). All images are composites of multiple focal planes (Helicon Focus software), with anterior to the left. The brackets in B, C’, D’, E’ and F represent the width of individual segments. Please note that there is a slight discordance in juvenile stages between the external segmentation (marked by the aforementioned brackets) and the internal segmentation of the nervous system (ganglia). The lateral region of the brain is indicated by asterisks (A-F), while the medial portion is designated by double open arrowheads (C-F) and the posterior region is highlighted with the double open arrow (D). *Ct-pax6* is expressed in the ventral nerve cord. At early and late stages, expression is restricted to a few cells (A, E’, F, filled arrowheads). Segmentally-repeated lateral diagonal stripes (B, C’, D’, black arrows) and medial spots (D’, white arrows) are distinguishable during the peak phase of *Ct-pax6* expression in the ventral nerve cord. *Ct-pax6* expression is visible outside the central nervous system in stage 7 larvae (D’, unfilled arrowheads) Abbreviations: pt: prototroch, st: stomodeum, tt: telotroch. Scale bars: 50 µm.

**Fig. 4.**
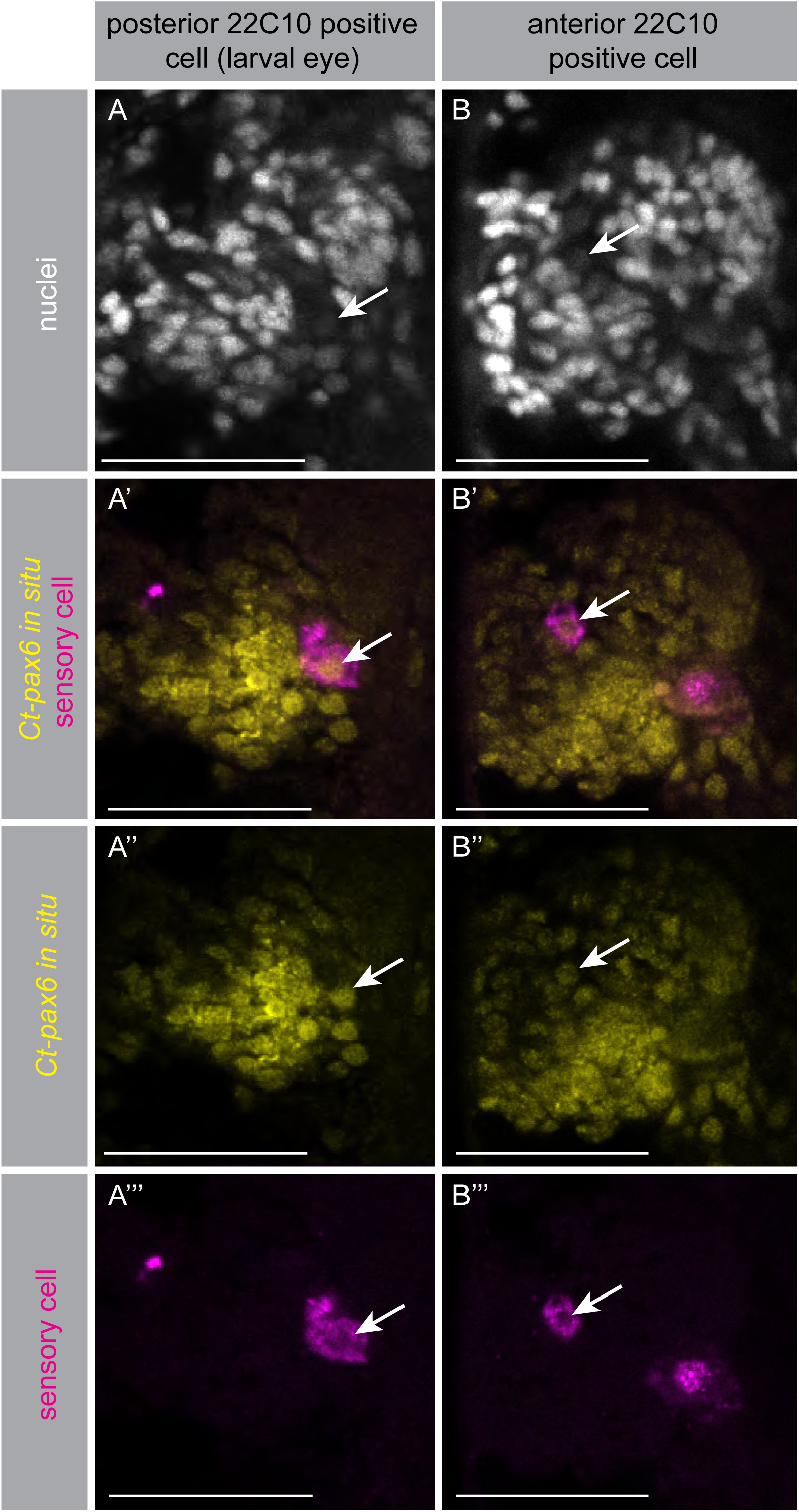
*Ct-pax6* expression in the eye of *C. teleta* larvae. Hoechst staining to label nuclei in white (nuclei; A, B), *Ct-pax6* whole mount *in situ* expression in yellow (*Ct-pax6*; A’, A’’, B’, B’’) and 22C10 antibody labeling of the sensory cell of the eye in magenta (sensory cells; A’, A’’’, B’, B’’’). A’ and B’ are merged channels of A’’ plus A’’’ and B’’ plus B’’’, respectively. The images shown are high magnification views of the brain-region of stage 6 larvae, with anterior to the left. Each image represents a single confocal slice only. Images from each column are from the same individual. A sub-region of both sensory cells expresses *Ct-pax6* (highlighted by arrows). Scale bars: 30 µm.

To examine if *Ct-pax6* was expressed in the larval photo-sensory cells, we combined *Ct-pax6 in situ* hybridization (yellow in Fig. 4A’, A’’, B’, B’’) and 22C10 antibody labeling (magenta in Fig. 4A’, A’’’, B’, B’’’), which specifically labels sensory cells associated with the eyes in *C. teleta* (see above). In larvae older than stage 5, there are two sensory cells on each side of the head, an anterior and a posterior sensory cell, and both have their cell bodies within the brain lobes (Fig. 4A, B, arrows, weakly stained). Many brain cells are *Ct-pax6*-positive (yellow in Fig. 4A’, A’’, B’, B’’), and *Ct-pax6* is also expressed within each of the sensory cells (magenta in Fig. 4A’, A’’’, B’, B’’’, arrows), indicating expression in the larval photoreceptors.

To summarize, *Ct-pax6* is expressed in distinct subdomains in the developing brain and ventral nerve cord of *C. teleta* larvae and juveniles. Additionally, both pairs of photo-sensory cells present in the larval head express *Ct-pax6*.

### Experimental strategy and morpholino validation

The pre-spliced *Ct-pax6* mRNA consists of ten exons and nine introns (Fig. 5A). This large number of exons found in the *Ct-pax6* gene is similar to what has been reported in various other species (Chisholm & Horvitz, 1995; Glardon, Callaerts, Halder, & Gehring, 1997; van Heyningen & Williamson, 2002). The paired box spans two exons, namely exon 3 and exon 4 (magenta in Fig. 5), while the homeobox spans three exons, namely exon 5, exon 6, and exon 7 (green in Fig. 5).

**Fig. 5.**
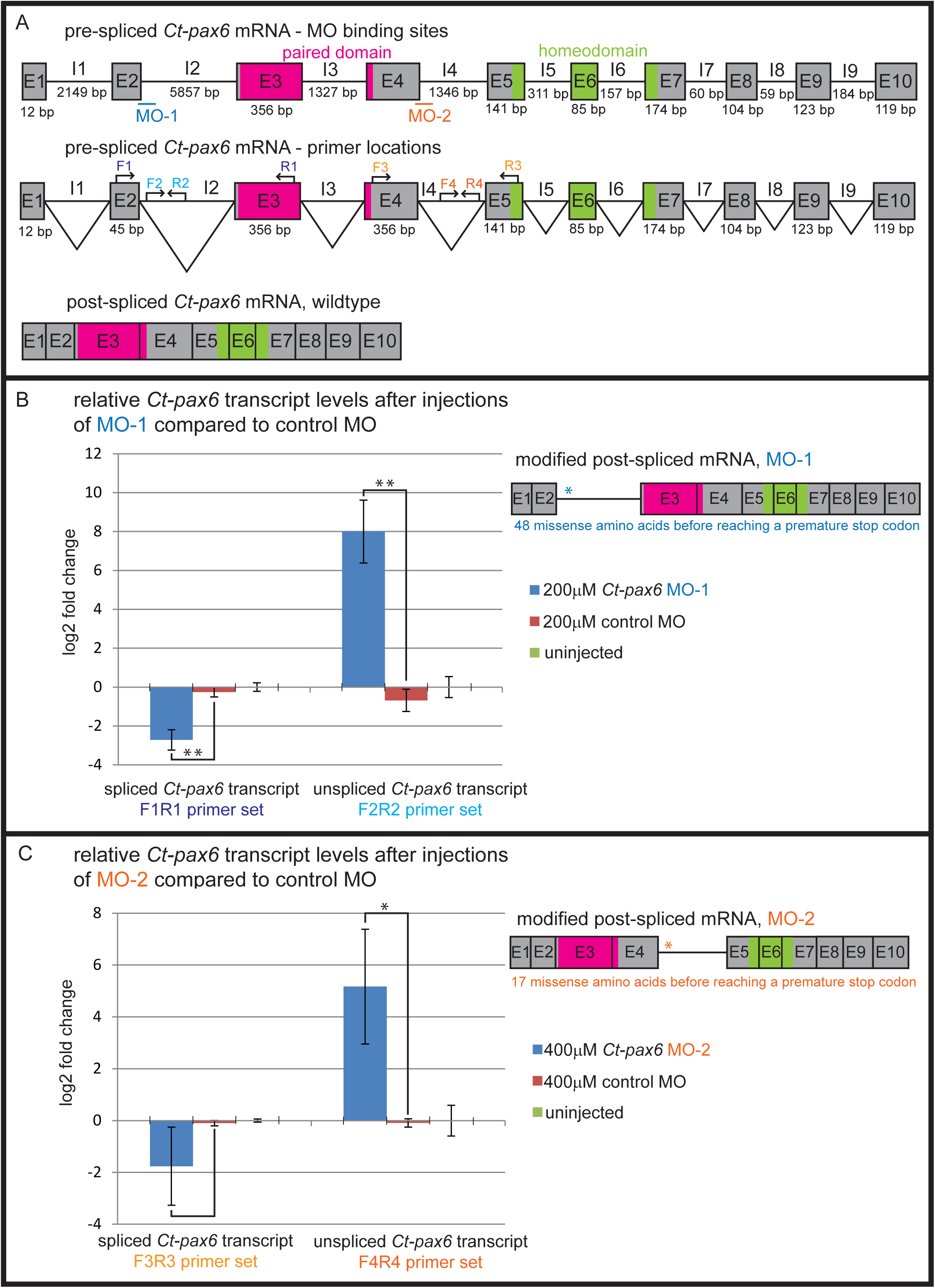
Experimental strategy and validation of the morpholino experiments. Schematic and graphic display of the morpholino (MO) strategy employed (A) and validation (B, C). Exons are represented by boxes that include the corresponding exon-number (E1 to E10). Introns are represented by lines with the corresponding intron-number above (I1 to I9). The size in base pairs (bp) of each exon and intron is shown underneath the box or line, respectively. The positions of the DNA-binding domains are marked, with the paired box in magenta and the homeobox in green. Elements associated with MO-1 are blue, while MO-2 associated elements are orange. (A) Morpholino strategy. The top schematic shows the organization of pre-spliced *Ct-pax6* mRNA with the location of the target sites of both morpholinos. MO-1 is at the second exon-intron-boundary and MO-2 is at the fourth exon-intron-boundary. The positions of the primer sets (F1&R1, F2&R2, F3&R3, and F4&R4) used to analyze the efficiency of splice blocking by the two MOs are indicated in the middle schematic. The bottom schematic depicts the wildtype post-spliced *Ct-pax6* mRNA. (B) Relative *Ct-pax6* transcript levels after injections of MO-1 (blue) compared to control MO (red). The bar graph shows the down-regulation of correctly spliced *Ct-pax6* transcript (left side) as well as the up-regulation of the unspliced *Ct- pax6* transcript (right side). The error bars on each column represent standard deviation. Note that a logarithmic scale is shown. Statistical significance where *p* = ≤ 0.01 is denoted by **, *p* values were calculated using a one-tailed Student’s *t*-test. The schematic on the right displays the modified post-spliced *Ct-pax6* mRNA after MO-1 injections, which results in the inclusion of the second intron. The asterisk marks the positions of premature stop codons included through modified splicing due to MO activity. (C) Relative *Ct-pax6* transcript levels after injections of MO-2 (blue) compared to control MO (red). The bar graph (logarithmic scale) shows the down- regulation of correctly spliced *pax6* transcript (left side) as well as the up-regulation of the unspliced *pax6* transcript (right side). The up-regulation is statistically significant: * ≤ 0.5; *p* value calculated using a one-tailed Student’s *t*-test. The schematic on the right displays the modified post-spliced *pax6* mRNA after MO-2 injections, which results in the inclusion of the fourth intron. The asterisk marks the positions of premature stop codons included through modified splicing due to MO activity.

For the morpholino (MO) knockdown experiments, two different splice-blocking morpholinos were designed to target the two different DNA-binding motifs of *Ct-pax6*, the paired domain and paired-type homeodomain. One morpholino, “MO-1”, binds to the second exon-intron-boundary (blue in Fig. 5A, top schematic). Sufficient splice-blocking activity of MO-1 results in the inclusion of the second intron (Fig. 5B, right schematic), which would introduce 48 missense amino acids before reaching a premature stop codon in the modified post-spliced mRNA (Fig. 5B, blue asterisk in right schematic). This means that if a protein is synthesized, it would contain neither a paired domain nor a homeodomain. In contrast, the second morpholino, “MO-2”, binds to the fourth exon-intron-boundary (orange in Fig. 5A, top schematic). Sufficient splice-blocking activity of MO-2 results in the inclusion of the fourth intron (Fig. 5C, right schematic), which would introduce 17 missense amino acids before reaching a premature stop codon in the modified post-spliced mRNA (Fig. 5C, orange asterisk in right schematic). In this case, the paired domain would be retained in the modified, truncated protein, but the DNA-binding homeodomain would be missing.

To check for potential off-target binding sites to mRNA sequences by the employed morpholinos, a Blast search against the publically available *Capitella* genome was performed. It is generally assumed that a five base pair mismatch spread throughout a 25-mer morpholino (typical length) results in a loss of knockdown activity. Every sequence with a significant alignment was investigated in detail since the activity of morpholinos are highly specific to either block translation (when targeted to the 5’ UTR or first 25 base pairs of coding sequence) or splicing (when targeted in introns near intron-exon boundaries. Morpholino homology to an off-target mRNA outside of these limited regions is unlikely to affect the expression of the off-target mRNA (Gene Tools). Taken all considerations into account, we found the control morpholino is very unlikely to bind and initiate a gene knockdown of any mRNA in *C. teleta*. Morpholinos specific to *Ct-pax6* show a 100% sequence homology with scaffold 29 only, which is the scaffold in which *pax6* is located. No other off-target mRNA binding sites that would affect gene expression could be found, including the potential second *Ct-pax6* gene, *pax6.2*, which is located on scaffold 38209.

RT-qPCR was used in order to validate the efficiency of both morpholinos. For each morpholino, two sets of specific primers were designed. One primer set was used to evaluate the level of spliced *Ct-pax6* transcript: F1&R1 for MO-1 and F3&R3 for MO-2 (Fig. 5A, middle schematic). A second primer set was used to examine the level of unspliced *Ct-pax6* transcript: F2&R2 for MO-1 and F4&R4 for MO-2 (Fig. 5A, middle schematic). Fold changes of relative *Ct-pax6* transcript levels were calculated (using *Ct-HPRT* as reference gene) and are displayed on a logarithmic scale in Fig. 5B and C. Three experimental treatment groups were compared: larvae resulting from *Ct-pax6* MO injections (blue bars in Fig. 5B, C), larvae resulting from control MO injections (red bars in Fig. 5B, C), and larvae resulting from uninjected embryos (light green bars in Fig. 5B, C). There are no statistically significant differences in *Ct-pax6* transcript levels between the control MO injected larvae and the uninjected larvae for all four primer pairs: F1&R1 with *p* = 0.26099, F2&R2 with *p* = 0.19828, F3&R3 with *p* = 0.29864, and F4&R4 with *p* = 0.58846 (two-tailed Student’s *t*-test). This shows that neither the injection process, nor the use of a generic standard control morpholino perturb the wildtype expression levels of *Ct-pax6* transcript in *C. teleta* larvae. There is a statistically significant down-regulation of correctly spliced *Ct-pax6* transcript in MO-1 injected larvae (*p* = 0.00462, one-tailed Student’s *t*-test) (Fig. 5B, left side of bar graph, F1&R1). Furthermore, the unspliced *Ct-pax6* transcript is statistical significantly up-regulated in MO-1 injected larvae (*p* = 0.00947, one-tailed Student’s *t*-test) (Fig. 5B, right side of bar graph, F2&R2). Larvae resulting from injections of MO-2 show a higher variability in *Ct-pax6* transcript levels than larvae injected with MO-1. Although there is no statistical significance (*p* = 0.07667, one-tailed Student’s *t*-test), a nearly 2-fold down-regulation of the correctly spliced *Ct-pax6* transcript in MO-2 injected larvae is seen when compared to larvae resulting from control MO injections (Fig. 5C, left side of bar graph, F3&R3). In contrast, a statistically significant up-regulation of the unspliced *Ct-pax6* transcript is detected in larvae resulting from MO-2 injections (*p* = 0.01317, one-tailed Student’s *t*-test) (Fig. 5C, right side of bar graph, F4&R4). Transcript could not be detected in any of the -RT-control samples (5 primer pairs tested: 4 experimental and the reference gene), demonstrating that there is no contamination with residual genomic DNA in any of the experimental samples. This control validates that the statistically significant up-regulation of both unspliced transcripts following MO injection is a real signal.

Phenotypic analysis of larvae resulting from injections with either morpholino resulted in a variety of different phenotypes that could be classified into five distinct categories (Fig. 6), with increasing severity levels ranging from one to five. Category I larvae exhibit a normal general morphology, but show a nervous system phenotype. Category II larvae are elongated but have malformations in their general morphology (such as a bent body axis, or disrupted segmentation) and also show a nervous system phenotype. Usually, the rate of development of *C. teleta* larvae is consistent within and between broods regarding the timing and appearance of characteristic morphological features (see summary above). In the case of category III larvae, injected embryos developed into larvae with many normal morphological features, but their development was delayed and morphological features were more similar to larvae that were reared for two days (stage 4 larvae) instead of the expected features at four days (compare Fig. 1A, E to Fig. 1C, G). In addition, most category III larvae also show malformations in their central and peripheral nervous system architecture. Injected animals that completed gastrulation, and then formed a ball of gastrulated cells with cilia, were designated category IV. Injected animals that did not complete early cleavages, but resulted in an arrested development, were defined as category V.

**Fig. 6.**
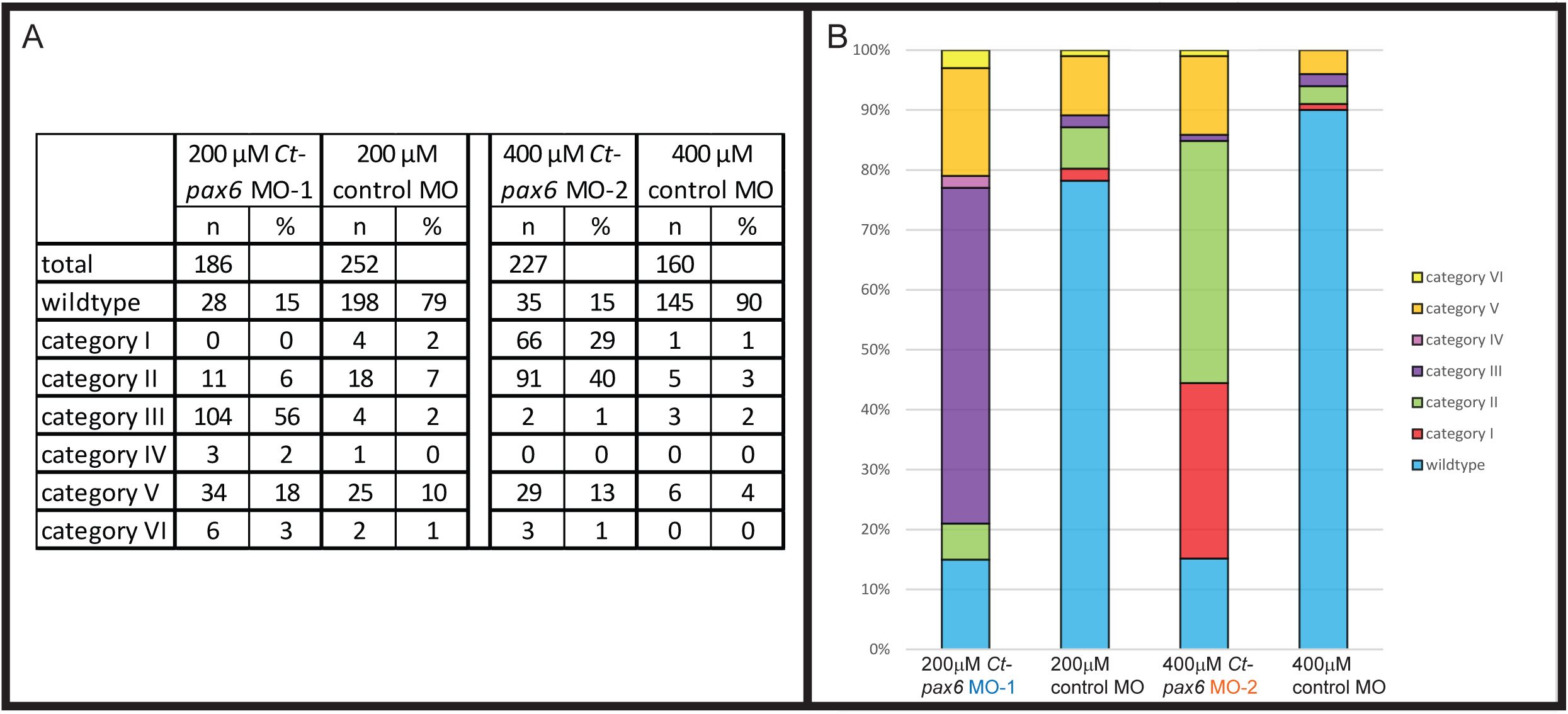
Summary of phenotypic characteristics after MO injections. Phenotypic analysis of larvae resulting from injections with either 200 μM MO-1 or 400 μM MO-2 morpholino resulted in a variety of different phenotypes which could be classified into five distinct categories. Category I larvae exhibit a normal general morphology, but show a nervous system phenotype. Category II larvae are elongated but have malformations in their general morphology and also show a nervous system phenotype. Category III larvae, exhibit a delayed development and mostly show malformations in their central and peripheral nervous system architecture. Injected animals that completed gastrulation, and then formed a ball of gastrulated cells with cilia, were designated category IV. Injected animals that did not complete early cleavages, but resulted in an arrested development, were defined as category V.

Because this is the first time we have used MOs to disrupt gene expression in *C. teleta*, we determined a suitable concentration for sufficient MO activity, including the control MO. In order to assess general toxicity of morpholinos on the development of *C. teleta* larvae, four increasing concentrations of the control MO were tested (Suppl. Fig. 2). The majority of 100 μM, 200 μM, 300μM, and 400 μM control MO injected embryos developed normally, resulting in wildtype phenotypic larvae (Suppl. Fig. 2, light blue portion of the bars). It can therefore be concluded, that injection of a morpholino into uncleaved *Capitella* embryos is not toxic as the general development of the embryos/larvae is not affected negatively. Judgment on the appropriate concentration of MO-1 was based on a morphological analysis. As for the control MO, four concentrations of MO-1 were tested to find a balance between toxicity (MO concentration too high) and insufficient MO activity (MO concentration too low): 100 μM, 200 μM, 300 μM, and 400 μM (Suppl. Fig. 3). A toxic effect was clearly seen for 300 μM and 400 μM MO-1 injections, as the majority of injected animals showed either a category IV or V phenotype (Suppl. Fig. 3). In contrast to injections of 400 μM and 300 μM, injections of 100 μM MO-1 resulted in a majority of larvae with a wildtype phenotype (Suppl. Fig. 3). 200 μM MO-1 was chosen as the appropriate MO-1 concentration for more detailed analyses due to a reasonable balance of resulting phenotypes (Fig. 6): a high proportion of larvae exhibited a category III phenotype (56%, *n* = 104/186), a low proportion of category V animals (3%, *n* = 6/186), and moderate proportion of either category IV animals (18%, *n* = 34/186) or wildtype larvae (15%, *n* = 28/186). The majority of the larvae resulting from injection of 200 μM control MO (74%, *n* = 198/252) developed into wildtype larvae (Fig. 6). The appropriate concentration of MO-2 was determined by analyzing levels of correctly spliced *Ct-pax6* transcript via RT-qPCR, using the primer set F3&R3 (Fig. 5A, middle schematic). Although injections of 200 μM MO-2 did not result in reduction of the correctly spliced *Ct-pax6* transcript (log2 fold change - 0.02389), a moderate down-regulation of the correctly spliced *Ct-pax6* transcript was achieved with a concentration of 300 μM MO-2 (log2 fold change −1.36907). Injections of 400 μM MO-2 was found to be suitable for the remaining experiments with an average log2 fold change of - 1.76044. 400 μM MO-2 injections into uncleaved eggs results in two prominent phenotypes: categories I (29%, *n* = 66/227) and II (40%, *n* = 91/227) (Fig. 6).

### Nervous system phenotype after *Ct-pax6* knockdown

Analysis of larvae resulting from injections with 200 μM MO-1 shows category III to be the predominate phenotype (Fig. 6; 56%, *n* = 104/186). This phenotype is only seen in larvae resulting from injections with 200 μM MO-1, but not in larvae resulting from injections with 200 μM control MO (Fig. 7, compare A, D, G to B, C, E, F, H, I). Category III larvae possess a number of morphological features typical of a stage 4 larva (Fig. 1A, E): a prototroch, telotroch, and stomodeum as well as segment anlagen in the ventral lateral region (Fig. 7B, C, open arrows). However, these larvae have not undergone elongation of the anterior-posterior body axis and lack all signs of segmentation in the trunk region. Antibodies against acetylated *a*-tubulin (Fig. 7D, E, F), serotonin (Fig. 7 G, H, I) and FMRFamide (data not shown) were used to assess the development of the nervous system in larvae resulting from 200 μM MO-1 injections (Fig. 7E, F, H, I), and compared to larvae resulting from 200 μM control MO injections (Fig. 7D, G). Although there is some variation in the phenotype (Fig. 7, compare B, E, H to C, F, I), typically there is a reduction in the number of neural fibers and neurons, and an overall disorganization of nerves. The few neural fibers and neurons are present in the brain (Fig. 7H, I, short arrows) and circumoral nerves (Fig. 7E, F, H, I, double open arrowheads). The main longitudinal connectives in the trunk are visible (Fig. 7E, F, H, I, long arrows), but display various malformations such as axon shortening (Fig. 7E, I), pathfinding errors (Fig. 7E, F), and presence of a single connective on only one side of the body (Fig. 7H, I). In some cases, nerves are severely mispositioned (Fig. 7E, F). There is also a lack of commissures in the trunk (compare Fig. 7D with E, F), which might be the result of missing ganglionic neurons, since no ganglia have formed. The anti- acetylated α-tubulin positive neurons in the head are missing (not shown). The peripheral nervous system is also affected in category III larvae. There is an absence of segmental nerves (Fig. 7F) and a reduced number of lateral longitudinal nerves in the trunk (Fig. 7E, F, double open arrows). To gain additional insight into the category III phenotype, one set of injected larvae was raised for a longer time period and the phenotype examined (6 days instead of 4 days, *n* = 201). Four days after injection, category III larvae (59.7 % of total injected (*n* = 120/201)) were separated from the group, and observed the following day. Of these, 69.16% (*n* = 83/120) of larvae had elongated but often retained malformations. The remaining 30.83% (*n* = 37/120) had elongated by day 6, but always displayed malformations, such as a bent shape body, absence of visible gut structures and a reduced pygidium. Therefore, 200 μM MO-1 injections resulted in a persistent phenotype that cannot be explained by a simple developmental delay, and there was a lack of recovery of the phenotype when animals were cultured for an extended period.

**Fig. 7.**
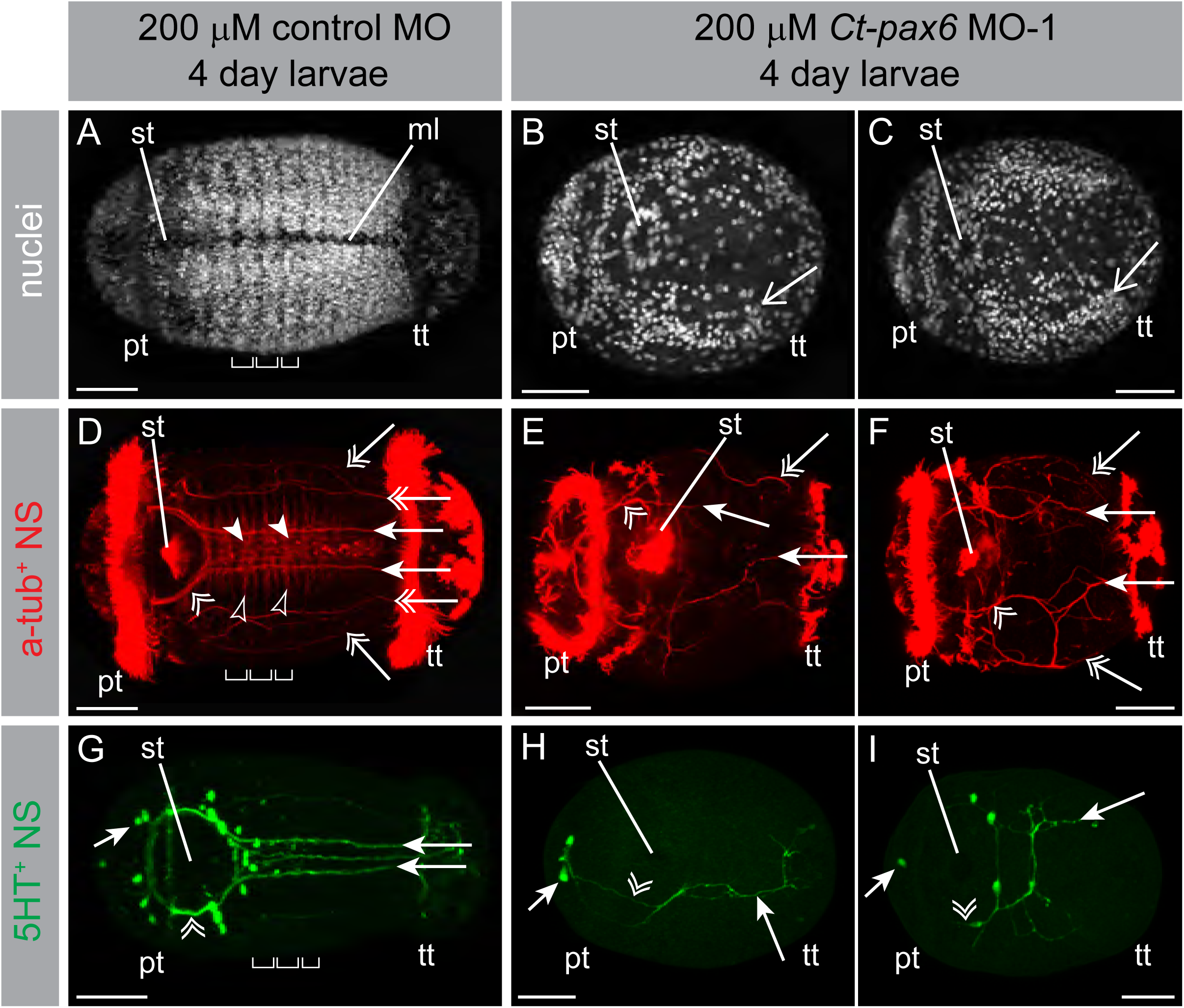
Phenotype of *C. teleta* larvae after MO-1 injections. Hoechst staining labels nuclei in white (nuclei; A-C), anti-acetylated α-tubulin labeling in red (a-tub^+^ NS; D-F) and anti-serotonin labeling in green (5HT^+^ NS; G-I). The images in A and D, B and E, and C and F are from the same individual. The developmental time for all larvae shown is four days. All images are z- stack projections, with anterior to the left. The open arrows in B and C show the location of the segment anlagen. The brackets in A, D and G represent the width of individual segments. The double open arrowheads in D–I point towards the circumoral nerves. The main connectives of the ventral nerve cord are indicated by the long arrows in D-I, while lateral longitudinal nerves are highlighted by double open arrows. The commissures (filled arrowheads) and selected individual segmental nerves (unfilled arrowheads) of the ventral nerve cord are highlighted in D. The short arrows in G and H mark the position of serotonin positive neurons in the brain. Abbreviations: ml: ventral midline, pt: prototroch, st: stomodeum, tt: telotroch. Scale bars: 50 µm.

To summarize, when reduction of *Ct-pax6* was achieved using a splice-blocking morpholino that results in a truncated protein lacking both the paired domain and the paired-type homeodomain, the majority of resulting larvae display a strong phenotype, category III, which has a highly reduced number of neurons and neural fibers. Many of the neurons and neural fibers present are disorganized, likely due to pathfinding defects.

Larvae resulting from injections with 400 μM of MO-2 also show a pronounced nervous system phenotype, that sharply contrasts with the normal phenotype (92%, *n* = 145/158) resulting from injections with 400 μM control MO (Fig. 8A, B, C). In larvae injected with 400 μM control MO, a nuclear stain shows segment boundaries and the position of the segmental tissue with respect to the ventral midline (Fig. 8A). Larvae resulting from 400 μM injections of MO-2 predominantly show either a category I or category II phenotype. First, category I larvae (29%, *n* = 66/227) exhibit a relatively normal gross morphology but exhibit abnormalities in nervous system development (Fig. 8D, E, F). They have visible segment boundaries, although the segmental tissue does not meet at the ventral midline as it normally would at this stage (Fig. 8D). Moreover, category I larvae have a reduced number of segmental nerves (Fig. 8E, unfilled arrowheads), commissures (Fig. 8E, filled arrowheads) and serotonin-positive neurons (Fig. 8F). Category II is characterized by larvae that are elongated, but show some malformations in their gross morphology and their nervous system architecture (40%, *n* = 91/227; Fig. 8G, H, I). Category II larvae exhibit highly disrupted trunk segmentation with few visible segment boundaries, and the medial edge of the presumptive segmental tissue shows a pronounced lateral displacement (Fig. 8G). The nervous system of category II larvae is under-developed. The main longitudinal connectives are thinner neural fibers and further apart than in control larvae (Fig. 8H, I). In addition, the distance between the nerves from the two contralateral sides increases as one moves posterior, instead of being parallel (Fig. 8H, I). There are few segmental nerves (Fig. 8H, unfilled arrowheads), a reduced number of serotonin-positive neurons in the brain-, circumoral-, and ventral nerve cord region (Fig. 8I), and a lack of commissures (Fig. 8H). Moreover, the lateral longitudinal peripheral nerves are reduced in number in category II larvae (Fig. 8H, double open arrows), but are present in category I larvae (Fig. 8E, double open arrows). The acetylated α- tubulin positive neurons that are typically located anterior to the brain are often altered by *Ct- pax6* transcript reduction (60%, *n* = 64/106). Wildtype larvae possess a bilateral pair with 3 neurons on each side (Fig. 1F, G, asterisks), while larvae resulting from injections with 400 μM MO-2 either have a reduced number of neurons (*n* = 39/106) (Fig. 8E, H, asterisks) or no neurons (*n* = 25/106). The asymmetric FMRF positive anterior mouth cell (F-AMC), which is located on the left side anterior to the mouth, is altered in 40% (*n* = 43/106) of larvae that result from 400 μM injections of MO-2. This is the only asymmetric, unpaired neuron identified in a previous study (Meyer et al., 2015), and the most common phenotype after *Ct-pax6* knockdown is a loss of the neuron (*n* = 38/106), but occasionally it is present and its position is shifted towards the midline (*n* = 5/106).

**Fig. 8.**
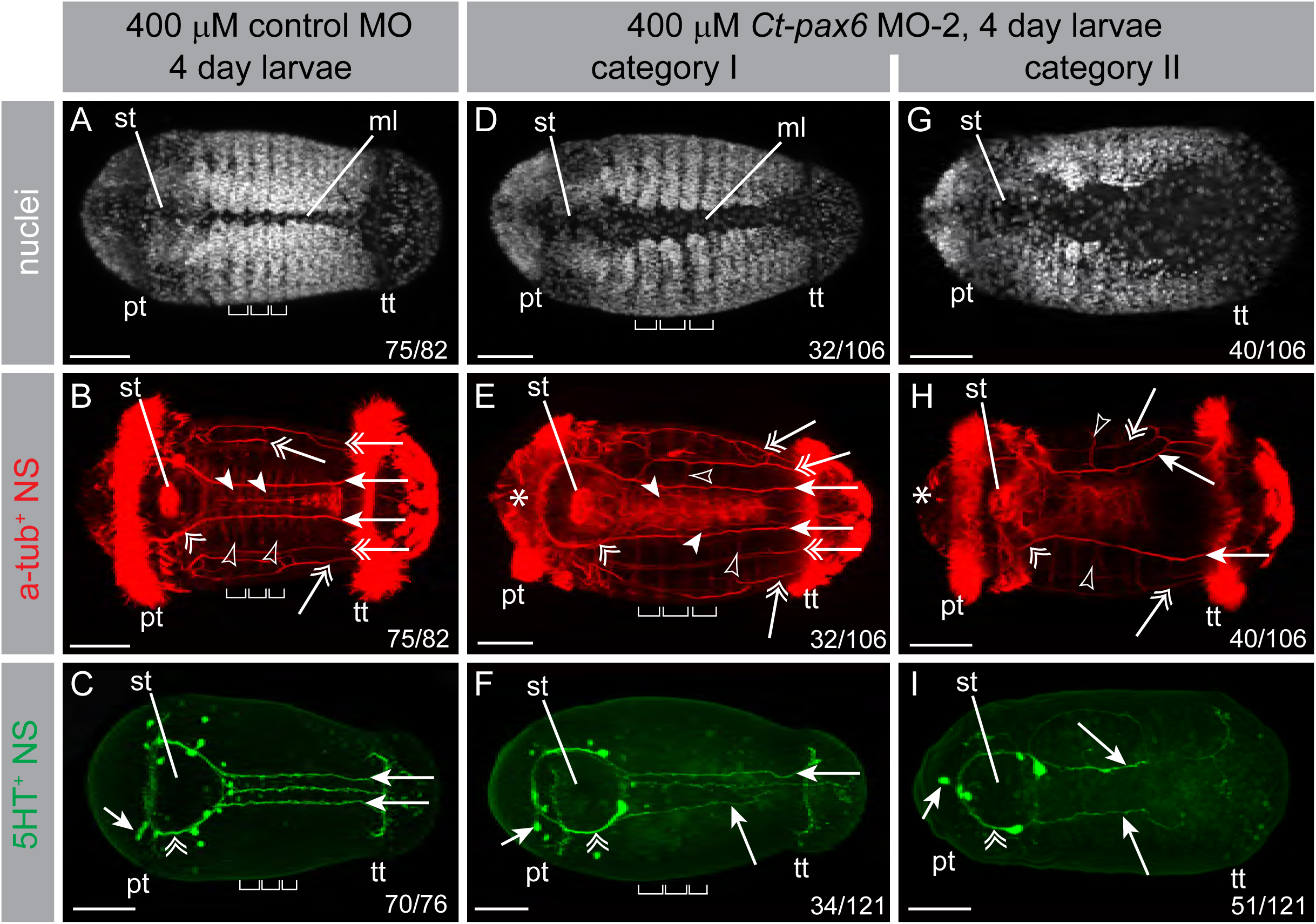
Nervous system phenotype of *C. teleta* larvae after MO-2 injections. Hoechst staining labels nuclei in white (nuclei; A, D, G), anti-acetylated α-tubulin labeling in red (a-tub^+^ NS; B, E, H) and anti-serotonin labeling in green (5HT^+^ NS; C, F, I). The images in A–B, D–E, and G– H are from the same individual. The developmental time for all larvae shown is four days. All images are z-stack projections, with anterior to the left. Numbers in the right bottom corner indicate the occurrence of the phenotype/sample size. The brackets in A–F represent the width of individual segments. The double open arrowheads in B, C, E, F, H, and I point towards the circumoral nerves. The main connectives of the ventral nerve cord are indicated by the long arrows in B, C, E, F, H, and I. The commissures in the ventral nerve cord are designated by filled arrowheads in B and E. Unfilled arrowheads highlight selected individual segmental nerves in B, E, and H, while double open arrows point towards the lateral longitudinal nerves. The asterisks in E and H mark the position of the anti-acetylated α-tubulin positive neurons in the head. The short arrows in C, F, and I show the position of serotonin positive neurons in the brain. Abbreviations: ml: ventral midline, pt: prototroch, st: stomodeum, tt: telotroch. Scale bars: 50 µm.

To summarize, when a morpholino is used that disrupts splicing of *Ct-pax6* resulting in a truncated protein that contains the paired domain, but does not contain the paired-type homeodomain, most of the resulting larvae show abnormalities in their nervous system. The predominate phenotypes resulting from injections of MO-2 (category I or category II) are considerably less severe than the predominant phenotype resulting from injections of MO-1 injections (category III, Fig. 6).

### Eye phenotype after *Ct-pax6* knockdown

Eye development was also analyzed in larvae resulting from *Ct-pax6* MO injections. The pigment cells and the sensory cells, two of the three cell types present in the eyes of *C. teleta*, were studied in detail in larvae resulting from injections with 200 μM MO-1 as well as 400 μM MO-2. Each of the two cell types was analyzed independently of each other. The severe phenotype resulting from 200 μM MO-1 injections is also visible with respect to eye development. None of the category III, IV, or V larvae had detectable pigment cells or sensory cells. In contrast, the relatively mild phenotypes resulting from injections of MO-2 (category I or category II) allowed for detailed analysis of the eye phenotypes (see below).

*C. teleta* larvae have a bilateral pair of pigment cells that are located dorsally at the lateral-posterior margin of the brain and in close proximity to the prototroch. This pigment cell position is evident in the majority (90%, *n* = 143/158) of larvae resulting from 400 μM control MO injections (Fig. 9A, double open arrowheads). In contrast, only 17% (*n* = 41/244) of larvae resulting from injections with 400 μM MO-2 have normal pigment cells. The majority of MO-2 injected larvae do not possess any pigment cells (40%, *n* = 97/244; Fig. 9B). Additionally, many larvae show abnormal pigment cell formation, such as the presence of only one pigment cell (Fig. 9C; 28%, *n* = 68/244), or a reduction in the size of the pigment cell (Fig. 9D; 16%, *n* = 38/244).

**Fig. 9.**
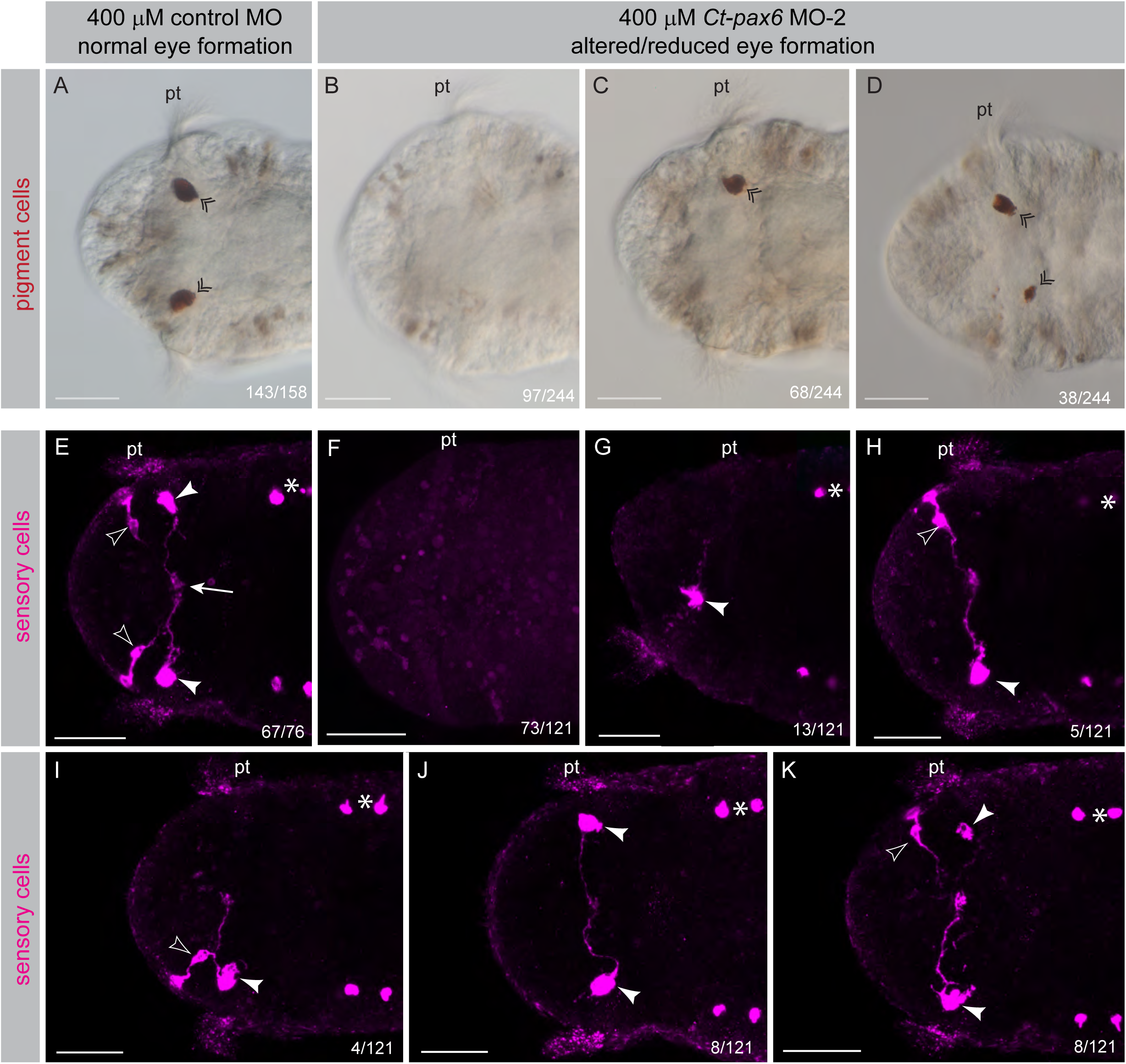
Eye phenotype of *C. teleta* larvae after MO-2 injections. Brightfield microscopic images to show the eye pigment in dark red (A-D) and 22C10 antibody labeling to label the sensory cells of the eye in magenta (E-K). The developmental time for all larvae shown is four to five days. All images are z-stack projections, with anterior to the left. Numbers in the right bottom corner indicate the occurrence of the phenotype/sample size. Double open arrowheads in A, C, and D mark the location of the pigment cell. The 22C10 antibody labels four sensory cells: a bilateral anterior pair (E, H, I, K unfilled arrowheads) and a bilateral posterior pair (E, G-K, filled arrowheads). The arrow in E marks the position in the midline of the brain where the axonal projections end. Segmentally iterated chaetae of the trunk are autofluorescent and therefore visible in some of the 22C10 images (asterisks). The posterior sensory cell (filled arrowheads) is associated with the pigment cell (double open arrowheads). Abbreviation: pt: prototroch

Formation of photo-sensory cells was investigated by reactivity with 22C10 antibody. Approximately 88% (*n* = 67/76) of larvae resulting from 400 μM control MO injections exhibit the wildtype arrangement of one anterior pair of sensory cells (Fig. 9E, unfilled arrowheads) and one posterior pair of sensory cells (Fig. 9E, filled arrowheads). Similar to the results for pigment cell formation, only a few larvae resulting from injections with 400 μM MO-2 show normal development of sensory cells (8%, *n* = 10/121). There are two predominate sensory cell phenotypes in larvae resulting from injections with 400 μM MO-2. The majority of larvae (60%, *n* = 73/121) do not possess any detectable sensory cells (Fig. 9F). Second, a substantial proportion of larvae (31%, *n* = 38/121) have an altered arrangement of sensory cells: presence of one sensory cell (Fig. 9G), presence of one anterior and one posterior sensory cell on different sides of the animal (Fig. 9H), sensory cells on one side of the animal only (Fig. 9I), presence of only anterior or posterior sensory cells on both sides of the head (Fig. 9J), or presence of three sensory cells (Fig. 9K). Both anterior and posterior sensory cells are affected in a similar proportion, which is in line with our finding that *Ct-pax6* is expressed in both cells (see above). It is noteworthy that category II larvae have no pigment (*n* = 91/91) or sensory cells (*n* = 52/52). To summarize, *Ct-pax6* reduction leads to larvae that either have no eyes or abnormal eye formation. This is evident for both the pigment cells and sensory cells.

### Downstream target analysis of *Ct-pax6*

Relative expression levels of selected putative target genes were analyzed for larval stages resulting from 400 μM MO-2 injections. Eight genes were selected for preliminary downstream target analysis of Ct-Pax6, and genes were chosen to reflect the observed phenotypes following reduction of *Ct-pax6* transcript, including loss of neural structures and the eye as well as reduced segmentation. First, the effect of Ct-Pax6 on three genes involved in neurogenesis was investigated: *neurogenin* (*Ct-ngn*), *neuroD* (*Ct-neuroD*), and *synaptotagmin 1* (*Ct-syt1*). *Ct-ngn* and *Ct-neuroD* show similar expression patterns during brain and ventral nerve cord neurogenesis in *C. teleta* larvae (Sur et al., 2017), and their temporal and spatial expression often coincides with *Ct-pax6* expression (M. K. personal observation), making both *Ct-ngn* and *Ct-neuroD* potential downstream target genes. *Ct-syt1* is a reliable marker of differentiated, mature neurons in *C. teleta* (Meyer et al., 2015), and its onset appears to be after *Ct-pax6*, which prompted us to investigate the effects of Ct-Pax6 knockdown on *Ct-syt1* expression levels. Second, we examined the effect of Ct-Pax6 on three genes belonging to the gene regulatory network for eye development: *six3/6* (*Ct-six3/6*), *r-opsin* (*Ct-r-opsin1*), and *eyes absent* (*Ct-eya*). Co-expression of *pax6* and *r-opsin* has been demonstrated in the annelid species *P. dumerilii* (Tessmar-Raible, Steinmetz, Snyman, Hassel, & Arendt, 2005). Moreover, *Ct-r-opsin1* coincides with 22C10 mAb labeling, which marks the photo-sensory cells in *C. teleta* larvae (see Fig. 2), making the gene an ideal target to test whether it is influenced by *Ct-pax6* transcript downregulation. Although there are no expression data for *Ct-six3/6* or *Ct-eya* in *C. teleta*, results from other species suggests that both genes have a conserved role in eye development (Halder et al., 1998; Mannini et al., 2004; Purcell et al., 2005). A third group of genes involved in segment boundary formation were analyzed, since a disruption of segmentation is a reoccurring phenotype of *Ct-pax6* downregulation (categories II and III). The segmentally iterated expression patterns of *engrailed* (*Ct-en*) (formerly *CapI-en*) (Seaver & Kaneshige, 2006) and a homologue of *lbx/ladybird* (E. C. S. personal observation) makes these genes good candidates for this purpose.

The average fold change values of four biological replicates (one biological replicate represents 150-200 pooled MO injected larvae) are displayed in the bar graph as Fig. 10. All three neurogenesis genes show mild relative transcript level reductions (−0.673, *Ct-neuroD*; −0.798, *Ct- syt1*; −0.939, *Ct-ngn*). A strong reduction of relative transcript levels is evident for the two eye genes *Ct-opsin* (−2.211) and *Ct-eya* (−2.179). Interestingly, *Ct-six3/6*, shows little reduction of relative transcript levels (−0.373). Likewise, there is little down-regulation of the segmentation genes with fold change values of 0.068 for *Ct-lbx* and −0.177 for *Ct-en*. None of these changes are statistically significant when *p* values were calculated using a one-tailed Student’s t-test. However, it is noteworthy that *p* values for the two segmentation genes are much higher (around 0.45) than the *p* values of the genes that are known to be *pax6* targets in other species (neurogenesis and eye genes with *p* values between 0.05 and 0.1). Nevertheless, our initial analysis of downstream targets of Ct-Pax6 indicate that in *C. teleta*, Ct-Pax6 has an inductive effect on *Ct-opsin* and *Ct-eya*, as well as a mild inductive effect on *Ct-ngn*, *Ct-neuroD*, and *Ct- syt1.*

**Fig. 10.**
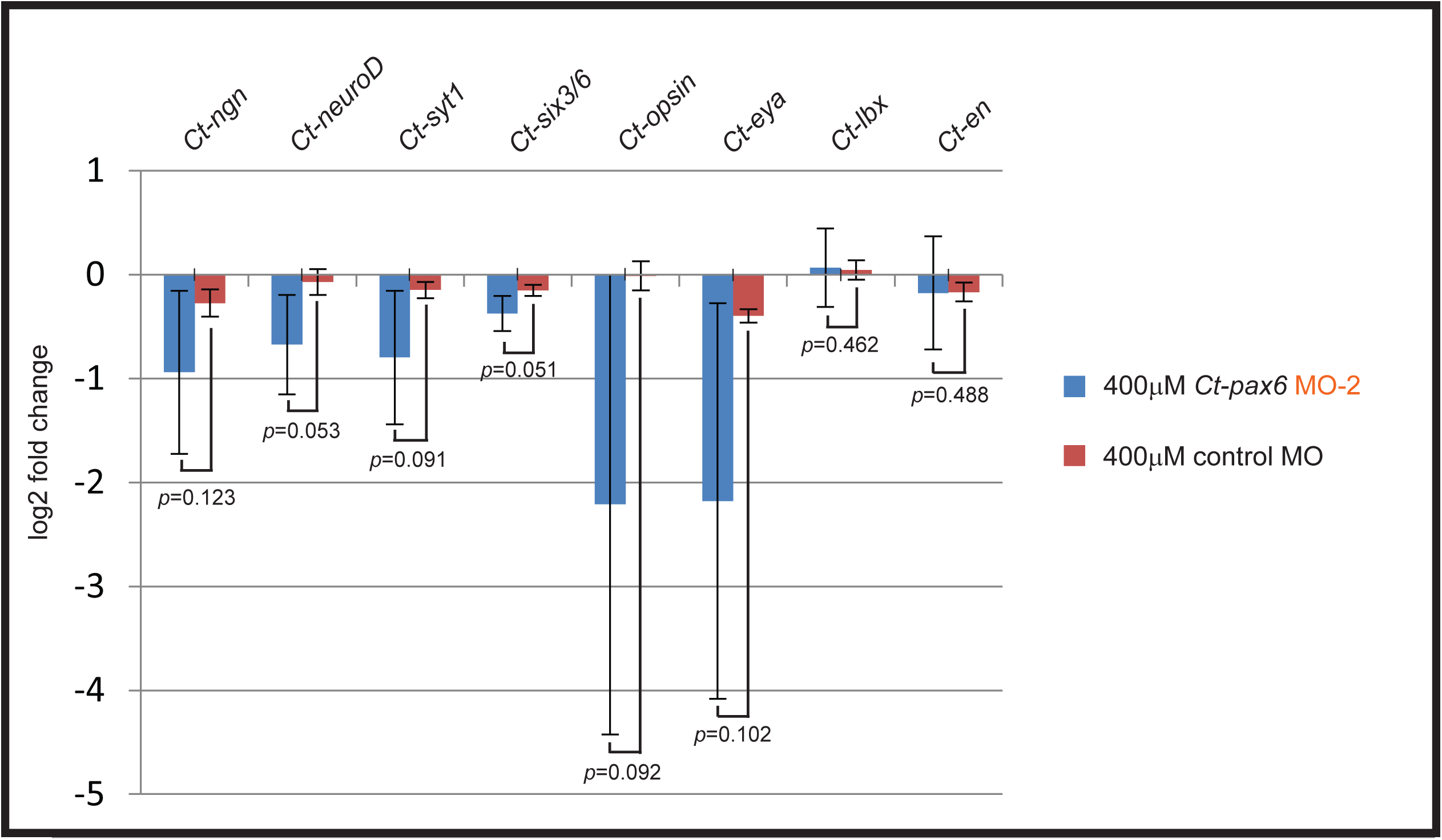
Relative transcript levels of selected *C. teleta* genes after MO-2 injections. Relative transcript levels after injections of MO-2 (blue) compared to control MO (red). The bar graph shows down-regulation of most transcripts. Error bars (representing standard deviation) as well as corresponding *p*-values are shown (calculated using a one-tailed Student’s *t*-test). Note that a logarithmic scale is shown. Abbreviations: *Ct-en*: *C. teleta engrailed*, *Ct-eya*: *C. teleta eyes absent*, *Ct-lbx*: *C. teleta ladybird*, *Ct-ngn*: *C. teleta neurogenin*, *Ct-syt1*: *C. teleta synaptotagmin 1*.

## Discussion

The present study represents the first publically available functional genomic study in *C. teleta*. Here, wildtype transcripts of *Ct-pax6* were reduced in larval stages using two different splice-blocking morpholinos. The results of this study provide both, evidence that morpholinos are efficient in *C. teleta* and result in specific phenotypes, as well as address the roles of both DNA-binding motifs of *pax6*.

### Evolutionary conservation of *pax6*

Gene orthology analysis previously identified a single *pax6* orthologue in the *C. teleta* genome (Seaver et al., 2012) that encodes the characteristic highly conserved DNA binding domains, a paired domain and a paired-type homeodomain, as well as a PST-rich transactivation domain. We do not address the second recently reported paired-less *pax6* orthologue (Zwarycz et al., 2015), since we could not experimentally validate transcript presence. As in *C. teleta*, the majority of species investigated so far possess a single *pax6* gene (defined by the presence of a complete paired box and paired-type homeobox), for example in the annelid *Platynereis dumerilii* (Arendt, Tessmar, de Campos-Baptista, Dorresteijn, & Wittbrodt, 2002), as well as in mouse (Hill et al., 1991) or human (Glaser, Walton, & Maas, 1992). There are some representatives, however, that have two complete *pax6* genes such as in *Drosophila melanogaster* (Czerny et al., 1999; Quiring, Walldorf, Kloter, & Gehring, 1994) and in the leech *Helobdella* (Quigley, Xie, & Shankland, 2007).

The genomic structure of the *Ct-pax6* gene includes ten exons that span over 12kb. *Pax6* genes from a number of different species are comprised of ten or more exons that occupy a large genomic region: human *pax6* with 14 exons over 22kb (Glaser et al., 1992; van Heyningen & Williamson, 2002), ascidian *pax6* with 10 exons (Glardon et al., 1997), nematode *pax6* with 14 exons (Chisholm & Horvitz, 1995), and mouse *pax6* with 13 exons (Kleinjan, Seawright, Childs, & van Heyningen, 2004). In *C. teleta*, the paired domain spans two exons, while the paired-type homeodomain is comprised of three exons. Both DNA-binding domains of Pax6 have been implicated in recognizing different downstream targets, and can have divergent roles during development (Ashery-Padan et al., 2000; Chi & Epstein, 2002; Haubst et al., 2004; van Heyningen & Williamson, 2002; Walcher et al., 2013).

### Role of *pax6* during eye development

The study of expression and functional involvement of the *pax6* gene during the specification and general development of the eye in various species has a long history (Noll 1993; Cvekl, & Callaerts, 2017). Expression of *pax6* associated with developing eyes has been demonstrated in most species investigated and is also found across major phylogenetic lineages: Lophotrochozoa, Ecdysozoa, and Deuterostomia. *C. teleta* belongs to the Annelida, a subclade of the Lophotrochozoa. In *C. teleta*, the larval eye consists of three different cell types, and *Ct-pax6* expression is only evident in the sensory cells, but not in the pigment or supporting cells. Association of *pax6* expression with the developing eye has been shown in a few other annelids, including the larval eye of *Platynereis dumerilii* (Arendt et al., 2002), and in the embryo of *Helobdella sp*. (Austin) (Quigley et al., 2007). Representatives of other lophotrochozoan subclades also express *pax6* in association with eye development, such as in squid (Hartmann et al., 2003; Tomarev et al., 1997), cuttlefish (Navet, Andouche, Baratte, & Bonnaud, 2009), ribbon worm (Loosli, Kmita-Cunisse, & Gehring, 1996), and plathyhelminthes (Pineda et al., 2002). The ecdysozoan lineage contains many examples where *pax6* expression is found in the developing eye (such as presence of transcripts in the optic lobes): in the onychophoran *Euperipatoides kanangrensis* (Eriksson, Samadi, & Schmid, 2013), the myriapod *Glomeris marginata* (Prpic, 2005), the wandering spider *Cupiennius salei* (Samadi, Schmid, & Eriksson, 2015), the fruit fly *Drosophila melanogaster* (Quiring et al., 1994), and the red flour beetle *Tribolium castaneum* (Yang et al., 2009). Even in the eyeless echinoderms, a *pax6* transcript is correlated with potential photo-sensory organs in the tube feet of sea urchins (Lesser, Carleton, Böttger, Barry, & Walker, 2011). There are many chordate representatives showing *pax6* transcripts expressed in the developing eye, including in *Branchiostoma floridae* (Glardon, Holland, Gehring, & Holland, 1998), lamprey (Murakami et al., 2001), lesser spotted dogfish (Ferreiro-Galve, Rodríguez-Moldes, & Candal, 2012), zebrafish (Nornes et al., 1998; Püschel, Gruss, & Westerfield, 1992), *Xenopus laevis* (Nakayama et al., 2015), and mouse (Walther & Gruss, 1991). Elaborate eyes are usually comprised of several different cell types and it is noteworthy that *pax6* expression is not necessarily detected in all of them. For example, in the blind Mexican cavefish, *Astyanax mexicanus*, *pax6* expression is reduced in the degenerating lens tissue but remains at normal levels in the retina and ciliary marginal zone (Strickler, Yamamoto, & Jeffery, 2001).

Although there are no functional studies investigating the role of *pax6* during eye development in species closely related to *C. teleta*, the reoccurring expression of *pax6* in the developing eye in a variety of animals (see above) suggests that *pax6* has a functional role in eye formation that is conserved across metazoans. Our *Ct-pax6* knockdown experiments show that normal eye development in *Capitella* larvae is highly impaired in MO injected animals. Interestingly, there is a high correlation between the absence of eyes and severe phenotypic alterations in nervous system development (categories II and III) with both MOs used in this study. The pronounced eye phenotype in *Capitella* larvae is in accordance with previous studies on a range of species and adds to a substantial body of evidence that the eye specification process relies on functional *pax6*, and is highly conserved across animals. Several individual lines of experimental evidence demonstrate this. First, ectopic expression of *pax6/ey* is sufficient for ectopic eye specification in *Drosophila* (Halder, Callaerts, & Gehring, 1995) as well as in *Xenopus* (Chow, Altmann, Lang, & Hemmati-Brivanlou, 1999). Moreover, *pax6* homologues from species as diverse as mouse, ascidian, squid, and cnidarian are capable of inducing ectopic eyes when ectopically expressed in *Drosophila* (Glardon et al., 1997; Halder et al., 1995; Kozmik et al., 2003; Tomarev et al., 1997). Although homozygous *pax6* mutants are typically lethal, heterozygous mutants display varying degrees of eye reduction or malformation, in humans (aniridia disease and Peter’s anomaly), mouse, rat, zebrafish, *Xenopus*, and *Drosophila* (Halder et al., 1998; Hill et al., 1991; Kleinjan et al., 2008; Matsuo et al., 1993; Nakayama et al., 2015). Thus, proper *pax6* function is essential for normal eye development in most animals investigated. However, there are some examples where eye development does not involve *pax6* expression, such as in the horseshoe crab (Blackburn et al., 2008).

To further explore the eye phenotype of *Ct-pax6* knockdown in larvae, three genes known to be important during eye development were analyzed using qPCR: *six3/6(optix)*, *r-opsin1*, and *eyes absent(eya)*. In *Drosophila*, *optix/six3/6* is a direct target of Ey/Pax6 (Ostrin et al., 2006), as is its mouse homologue *six3/6* (Purcell et al., 2005), and *six3/6* is predicted to be a Pax6 target in zebrafish (Coutinho et al., 2011). Expression of *six3/6* in the developing eye is widespread among ecdysozoans and deuterostomes (Kumar, 2009b). Direct regulation of *rhodopsin1* by Ey/Pax6 has been demonstrated for *D. melanogaster* (Sheng et al., 1997). It has been confirmed that *eya* is a direct target of Ey/Pax6 in *Drosophila* (Halder et al., 1998; Ostrin et al., 2006) and mouse (Purcell et al., 2005), and *eya* is predicted to be a Pax6 downstream target in zebrafish (Coutinho et al., 2011). Although the result is not statistically significant (due to high biological variability), there is a strong reduction of relative transcript levels for *Ct-opsin* and *Ct-eya* in *Ct- pax6* knockdown larvae. This hints at an evolutionary conservation of *opsin* and *eya* as downstream targets of Pax6. *Ct-six3/6*, however, only shows a minimal reduction of relative transcript levels. Previous studies of *six3/6* function have mainly been conducted in arthropod or vertebrate representatives, where a functional involvement in lens formation has been demonstrated. Many lophotrochozoan eyes do not undergo lens development, which could be a reason why there is no available expression data (Pineda & Saló, 2002; Steinmetz et al., 2010). Indeed, a comparative RNA-sequence study of developing eyes of *Nautilus* and pygmy squid revealed that *six3/6* and many downstream targets are not expressed in *Nautilus* (Ogura et al., 2013). The authors suggest that changes in the *six3/6* pathway might have led to the evolution of pinhole eyes in *Nautilus* (Ogura et al., 2013). The pinhole eye does not contain a lens and is a highly derived version of cup-like eyes, which are very common amongst lophotrochozoans. In contrast, all coleoid cephalopods (including pygmy squid) possess highly derived camera lens eyes. In summary, our qPCR data suggest that there may be conservation of an eye gene regulatory network that contains *pax6* and *eya* during eye determination, and *opsin* during later stages of eye development. Although further analysis is necessary to explore this in detail, these findings are consistent with the proposed retinal gene network of *D. melanogaster* (Cvekl & Callaerts, 2017; Kumar, 2009a).

### Role of *pax6* during nervous system development

In *C. teleta* larvae, initial *Ct-pax6* expression coincides with the onset of the development of the central nervous system. Further maturation of the central nervous system is accompanied by expansion of *Ct-pax6* expression, in distinct subpopulations in the brain and in a segmentally iterative pattern in the ventral nerve cord. These distinct subpopulations indicate a role for *Ct- pax6* in neuronal subtype specification in *C. teleta*, as is seen in mouse and *Xenopus* (Aleen Remez et al., 2017; Dulcis et al., 2017). The subsequent restriction to a reduced number of *Ct- pax6* positive cells in *C. teleta* larvae, could also indicate a role in later, newly forming neurons. A similar expression pattern of *pax6* in the central nervous system during embryonic/larval development is found in all species investigated thus far: typically, expression is in subdomains in the brain and there is an iterative pattern of a subset of cells in the (ventral) nerve cord. Within lophotrochozoans, examples include: planarians (Pineda et al., 2002), leech (Quigley et al., 2007), the annelid *P. dumerilii* (Denes et al., 2007), the ribbon worm (Loosli et al., 1996), and the mollusc *Wirenia argentea* (Scherholz et al., 2017). Within ecdysozoans, examples include *E. kanangrensis* (Eriksson et al., 2013), *Limulus polyphemus* (Blackburn et al., 2008), a myriapod (Prpic, 2005), *T. castaneum* (Yang et al., 2009) and *D. melanogaster* (Quiring et al., 1994), and within chordates the tunicate, *Phallusia mammillata* (Glardon et al., 1997), and the lamprey (Murakami et al., 2001).

In *C. teleta*, expression of *Ct-pax6* transcript suggests that it has at least two roles during neurogenesis. First, it has a role in early neural specification and differentiation, which is represented by the broad *Ct-pax6* expression during larval stages 4-7, and a later role in specifying different neuronal subtypes. A recent paper on spatiotemporal regulation of various genes involved in the development of *C. teleta* found that genes such as *soxB1*, *soxB* and *neurogenin* may act to keep the neural precursor cell in a proliferative state (Sur et al., 2017). *Ct- pax6* is expressed in similar stages to each of these three genes, but in a smaller number of cells during larval stages 4 and 5 (neural specification). In contrast, *Ct-pax6* has a much broader expression pattern during larval stages 6 and 7 (neural differentiation). Furthermore, the authors propose that *musashi* and *neuroD* are involved in neuronal differentiation with a peak expression in larval stage 6 (Sur et al., 2017). *Ct-pax6* expression is evident much earlier than these two genes, but is partially overlapping with both patterns during larval stage 6. This adds further evidence that *Ct-pax6* transcripts are active during neural precursor maintenance as well as neural differentiation. This is in accordance with previous studies that have shown *pax6* to act during different stages of neurogenesis (Blake & Ziman, 2014; Sansom et al., 2009; Shaham, Menuchin, Farhy, & Ashery-Padan, 2012). A later role in specifying different neuronal subtypes, including those of late born neurons (since not all neurons are born at the same time), is suggested by the spatially restricted *Ct-pax6* expression pattern in the brain and ventral nerve cord of larval stages 8-9 and juvenile worms. Studies in other animals support the hypothesis that *pax6* has important functions during neuron subtype specification (Aleen Remez et al., 2017; Dulcis et al., 2017; Kroll & O’Leary, 2005; Philips et al., 2005).

Microinjection of either MO-1 or MO-2 demonstrates that down-regulation of the wildtype *Ct- pax6* transcript and an increase in incorrectly spliced transcript in *C. teleta* larvae, disrupts normal nervous system development. Phenotypic changes include a reduced number of neural fibers and neurons in the brain and the ventral nerve cord. In mouse, it has been shown that Pax6 function is important for neural stem cell renewal (Sansom et al., 2009), and mouse homozygous mutants show a premature cessation of neural progenitor proliferation resulting in fewer neural stem cells present to differentiate into neurons (Philips et al., 2005). This is in line with our MO results, where fewer neurons are born. Specifically, the knockdown of *Ct-pax6* transcript with both DNA binding motifs missing (MO-1) results in larvae with a highly reduced number of neurons and neural fibers that form a rudimentary nervous system (anlagen of the brain, circumoral nerves and main longitudinal connectives), resulting in a lack of cells for further maturation. This seems to result in a developmental delay of the entire larvae. Similarly, zebrafish that were injected with a *pax6b* morpholino or simultaneous injections of *pax6* morpholinos targeting both *pax6* homologs, not only show disrupted eye development, but also a general developmental delay (Kleinjan et al., 2008). In *C. teleta*, along with a decreased number of neurons/neural fibers, there is a general disorganization of the neural fibers, consistent with pathfinding defects. For example, the longitudinal connectives are positioned too far apart. These nerves are typically positioned at the margin of the segment anlagen and ventral midline, which might be explained by the presence of pathfinding cues at the margin of the segment anlagen. *Pax6* mutants in *Drosophila* and mouse also show axon pathfinding defects (Jones, López-Bendito, Gruss, Stoykova, & Molnár, 2002; Mastick, Davis, Andrew, & Easter, 1997; Noveen, Daniel, & Hartenstein, 2000), and these phenotypes have been interpreted as being due to defects in axon guidance. Moreover, *ey/pax6 Drosophila* mutants have altered brain morphologies (in particular in the domains associated with vision and olfaction), and display uncoordinated locomotion (Callaerts et al., 2001). Similarly, during murine forebrain development, *pax6* is essential for cell proliferation, fate and patterning as well as for the development of the olfactory system (Haubst et al., 2004; Nomura et al., 2007; Walcher et al., 2013). Thus, *pax6* function is crucial for nervous system development in a range of animals, including in neurogenesis.

To gain further insight into the nervous system phenotype of *Ct-pax6* knockdown larvae, we analyzed three genes involved in neurogenesis via qPCR: *neurogenin* (*Ct-ngn*), *neuroD*, and *synaptotagmin 1* (*Ct-syt1*). Both, *ngn* and *neuroD* are direct targets of Pax6 in mouse and show reduced expression levels in *pax6*-mutant mice (Holm et al., 2007; Sansom et al., 2009; Visel et al., 2007). *SytIV* is predicted to be a direct target of Ey/Pax6 in *D. melanogaster* (Ostrin et al., 2006). Due to variation across biological replicates in our experiments, there is no statistically significant reduction, but nevertheless there is a slight reduction of transcription levels of all three neurogenesis genes in *Ct-pax6* knockdown larvae. Taken together with the observations that these three genes are targets of Pax6 in other species, one might speculate that *ngn*, *neuroD* and *sytIV* are conserved downstream targets of Pax6. Since less attention has been given to the roles of *pax6* during central nervous system development, future functional studies will further unravel its exact functions.

We also analyzed two genes involved in segment boundary formation, *engrailed* and *ladybird*, since disruption of trunk segmentation is a reoccurring phenotype of both *Ct-pax6* morpholinos used in this study. *Lbx* was chosen based on data from another annelid species *Platynereis* (Saudemont et al., 2008), in which *lbx/ladybird* expression suggested to play a role in segment boundary formation. Pax6 has been linked to boundary formation in the mouse brain, and it has been demonstrated that *en* is repressed by Pax6 to form the di-mesencephalic boundary (Mastick et al., 1997; Matsunaga et al., 2000). We found, however, that there is no significant downregulation of *Ct-en* or *Ct-lbx*. Nevertheless, Ct-Pax6 has a strong influence on segment boundary formation in *Capitella* larvae, potentially acting through a currently unknown segmentation pathway.

### Conserved functional domains of Pax6 and their roles

Employment of two different morpholinos to achieve functional knockdown of *Ct-pax6* transcripts results in larvae with distinct phenotypes. This is expected since the morpholinos were designed to block splicing at different sites in the *pax6* transcript and hence to affect translation of the two conserved DNA binding domains differently. MO-1 resulted in a transcript lacking both the paired box and the paired-type homeobox, while MO-2 resulted in a transcript lacking only the paired-type homeobox. The comparison of the different phenotypes for both MOs suggests that in *C. teleta* the paired domain and the paired-type homeodomain of Ct-Pax6 might have different downstream DNA-binding targets, which has been demonstrated for other species (Chi & Epstein, 2002; Haubst et al., 2004; van Heyningen & Williamson, 2002; Walcher et al., 2013).

### The homeodomain of Pax6 is involved in eye development

Both MOs used in this study result in larvae with pronounced eye developmental phenotypes. Although a detailed analysis of larvae lacking both functional Pax6 domains (resulting from injection of MO-1) is difficult due to the phenotypic severity of the resulting larvae, the majority had neither pigment nor sensory cells (over 80%). If only the DNA binding activities of the paired-type homeodomain is diminished (resulting from injections of MO-2), eye development in the resulting larvae was highly disrupted; ranging from a complete lack of eye cells, a reduction in size, to abnormal development. These features affected the pigment cells (84%) as well as the sensory cells (91%). Our results are in line with previous studies, which have shown that the paired-type homeodomain of Pax6 is crucial for eye development (Ashery-Padan et al., 2000; van Heyningen & Williamson, 2002). There are a couple of studies of mutants that impair the homeodomain only, for example the original *small eye* mutants in mouse and rat or the hypomorph mouse mutant *Pax64Neu* (Favor et al., 2001; Hill et al., 1991; Matsuo et al., 1993). Typically, homozygous mouse mutants lack eyes and are embryonic lethal, while heterozygous littermates display abnormal eye development including reduction in eye size. Furthermore, Nakayama et al. (2015) showed that targeted mutagenesis in *Xenopus* results in froglets with abnormal eye development when the homeobox of *pax6* is removed. The authors note however, that homozygous *pax6 Xenopus* mutants do not lose the complete eye structure as seen in mouse or rat (Nakayama et al., 2015). Therefore, mutations in the paired-type homeobox of *pax6* only are sufficient to disrupt normal eye development.

### The paired domain of Pax6 is involved in neurogenesis

The paired domain has been implicated with multiple roles during neurogenesis, such as neuronal subtype specification, axon guidance and neuronal proliferation (Haubst et al., 2004; Huettl et al., 2016; Walcher et al., 2013). However, these effects are variable across species, and suggest that the evolutionary functions of the paired domain are less conserved. Moreover, it has been shown that the paired-type homeodomain is also involved in neuronal regulation, for example in the boundary formation of the mouse brain (Haubst et al., 2004). In *C. teleta*, both DNA binding domains seem to be involved in regulating neurogenesis, since both morpholinos result in larvae with a reduced number of neurons and neuronal fibers.

#### Summary and concluding Remarks

Functional studies using splice-blocking morpholinos in the marine annelid *C. teleta* sufficiently reduce wildtype transcript levels of target genes, and are highly reproducible. Transcript expression studies in *C. teleta* larvae and juveniles confirm the presence of *Ct-pax6* in discrete subdomains of the brain and ventral nerve cord as well as in the sensory cells of the larval eye. Reduction of *Ct-pax6* transcript levels results in larvae with both severe eye and nervous system developmental defects. It is notable that from our analysis, we did not identify a simple segregation of function between a role in eye development and a role in neurogenesis between the paired domain and the homeodomain of Pax6 in *C. teleta*. Further studies by targeted disruption of the paired domain, while leaving the paired-type homeodomain intact and analyzing larval phenotypes for defects in eye formation and in the nervous system may provide insights into functional differences of these two domains. It has recently been proposed that the ancestral role for *pax6* maybe have been is in head patterning and neurogenesis, and that acquiring a role in eye development occurred secondarily (Cvekl and Callaerts, 2017). Taken together, our data supports the concept of an evolutionarily conserved role for *pax6* in the development of both the nervous system and eyes.

## Acknowledgements

The authors thank Dr. Danielle de Jong, Alexis Lanza and Leah Dannenberg for helpful suggestions and comments regarding the manuscript. The authors would like to thank Dr. Mansi Srivastava for sharing a protocol on the production of self-made TSA. This work was supported by the National Science Foundation (IOS1457102 to ECS).

---

**Supplementary Fig. 1. Summary of phenotypic characteristics after various MO-1 injections.** Phenotypic analysis of larvae resulting from injections with 100 μM, 200 μM, 300 μM or 400 μM MO-1 morpholino resulted in a variety of different phenotypes which could be classified into five distinct categories. Category I larvae exhibit a normal general morphology, but show a nervous system phenotype. Category II larvae are elongated but have malformations in their general morphology and also show a nervous system phenotype. Category III larvae, exhibit a delayed development and mostly show malformations in their central and peripheral nervous system architecture. Injected animals that completed gastrulation, and then formed a ball of gastrulated cells with cilia, were designated category IV. Injected animals that did not complete early cleavages, but resulted in an arrested development, were defined as category V.

**Supplementary Table 1. Summary of primers employed in this study.** The third and fourth column, location and abbr., apply to the MO validation primers only and refer to Fig. 5A.

